# The enhanced multi-tissue atlas of regulatory effects in cattle

**DOI:** 10.64898/2026.03.18.712441

**Authors:** Houcheng Li, Huicong Zhang, Di Zhu, Pengju Zhao, Zhenyu Wei, Jingsheng Lu, Mian Gong, Qi Zhang, Weijie Zheng, Xinfeng Liu, Dailu Guan, Jinyan Teng, Qin Lin, Yongjie Tang, Yahui Gao, Shanjiang Zhao, Zhe Zhang, Junting Du, Chao Fang, Bingxing An, Bingjin Lin, Haihan Zhang, Min Tian, Jingjing Tian, Siqian Chen, Wansheng Liu, Yanan Wang, Mingshan Wang, Eveline M. Ibeagha-Awemu, Richard P. M. A. Crooijmans, Martijn F. L. Derks, Marta Gòdia, Ole Madsen, Hubert Pausch, Alexander S. Leonard, Laurent Frantz, David E. MacHugh, John F. O’Grady, Iuliana Ionita-Laza, Xin Zhao, Leluo Guan, Huaijun Zhou, Emilio Mármol-Sánchez, Monique G. P. van der Wijst, Xubin Lu, Hui Jiang, Zhangping Yang, Qien Yang, Qinyou Liu, Chuang Xu, Moli Li, Yali Hou, Zhangyaun Pan, Yan Chen, Ruidong Xiang, Mathew Littlejohn, Emily L. Clark, Wenfa Lyu, Yuwen Liu, Lin Jiang, Peng Su, Xuexue Liu, Senlin Zhu, Minghui Jia, Huizeng Sun, Bo Han, Yi Zhang, Ying Yu, Dongxiao Sun, Yaokun Li, Dewu Liu, Goutam Sahana, Zexi Cai, Mogens Sandø Lund, John B. Cole, Li Ma, Jicai Jiang, Wenjian Li, Yang Wu, Jianbin Li, Jun Teng, Qin Zhang, Xiao Wang, Xuemei Lu, Yu Jiang, Yang Zhou, Yu Wang, Bingjie Li, Peter Sørensen, Lingzhao Fang

**Author notes:** Co-corresponding authors: Xiao Wang; Xuemei Lu; Yu Jiang; Yang Zhou; Yu Wang; Bingjie Li; Peter Sorenson; Lingzhao Fang. Co-first authors (Houcheng Li, Huicong Zhang, Di Zhu, Pengju Zhao, Zhenyu Wei, Jingsheng Lu). Emails for all authors: Houcheng Li; Huicong Zhang; Di Zhu; Pengju Zhao; Zhenyu Wei; Jingsheng Lu; Mian Gong; Qi Zhang; Weijie Zheng; Xinfeng Liu; Dailu Guan; Jinyan Teng; Qin Lin; Yongjie Tang; Yahui Gao; Shanjiang Zhao; Zhe Zhang; Junting Du; Chao Fang; Bingxing An; Bingjin Lin; Haihan Zhang; Min Tian; Jingjing Tian; Siqian Chen; Wansheng Liu; Yanan Wang; Mingshan Wang; Eveline M. Ibeagha-Awemu; Richard P. M. A. Crooijmans; Martijn F. L. Derks; Marta Gòdia; Ole Madsen; Hubert Pausch; Alexander S. Leonard; Laurent Frantz; David E. MacHugh; John F. O’Grady; Iuliana Ionita-Laza; Xin Zhao; Leluo Guan; Huaijun Zhou; Emilio Mármol Sánchez; Monique G. P. van der Wijst; Xubin Lu; Hui Jiang; Zhangping Yang; Qien Yang, Qinyou Liu; Chuang Xu; Moli Li; Yali Hou; Zhangyuan Pan; Yan Chen; Ruidong Xiang; Mathew Littlejohn; Emily L. Clark; Wenfa Lyu; Yuwen Liu; Lin Jiang; Peng Su; Xuexue Liu; Senlin Zhu; Minghui Jia; Huizeng Sun; Bo Han; Yi Zhang; Ying Yu; Dongxiao Sun; Yaokun Li; Dewu Liu; Goutam Sahana; Zexi Cai; Mogens Sandø Lund; John B. Cole; Li Ma; Jicai Jiang; Wenjian Li; Yang Wu; Jianbin Li; Jun Teng; Qin Zhang.

## Abstract

Cattle are integral to global food security, yet the molecular architecture of their complex traits remains poorly understood. Here, we present the **Cattle Genotype–Tissue Expression (CattleGTEx) Phase 1** resource (https://cattlegtex.farmgtex.org/), a substantial expansion of the pilot study. By leveraging 12,422 RNA-seq profiles across 43 tissues and 82 breeds, we characterized 433,972 primary and 161,428 non-primary regulatory effects spanning seven molecular phenotypes. This high-resolution atlas resolves 75% of GWAS signals for 44 complex traits, significantly addressing the "missing regulation" in livestock. We propose a genetic regulatory model demonstrating how variants across multiple biological layers interact with specific biological contexts to shape phenotypic variation. Furthermore, CattleGTEx elucidates mechanisms underlying adaptive evolution between *Bos taurus* and *Bos indicus*, as well as artificial selection in dairy and beef breeds. Finally, by mapping evolutionary constraints on these regulatory effects, we demonstrate the translational value of this resource for prioritizing causal variants in human complex diseases. Together, Phase 1 of CattleGTEx provides a transformative framework for functional genomics, precision breeding, and comparative genetics.

## Introduction

Since their domestication approximately 10,000-8,000 years ago, cattle have played a crucial role in human societal development and global food security, providing high-quality animal protein, labour, economic value, and cultural significance^1^. As they spread across the globe alongside human migration, cattle adapted genetically to diverse local environments ranging from cold, high-altitude mountains (e.g., Tibetan cattle) to tropical climates (e.g., Zebu)^2^. Modern breeding further intensified selection for production-related traits, resulting in specialized breeds with distinct physiological characteristics, such as the Holstein for dairy and Angus for beef^3^. Today, rising demand for sustainable food systems places new environmental and welfare pressures on cattle industry, necessitating improved production efficiency via precision breeding^4–6^. Beyond agriculture, the substantial genetic similarities between cattle and humans, particularly in immune, metabolic, and reproductive pathways, position cattle as valuable human biomedical models. These include conditions such as polycystic ovarian syndrome (PCOS), tuberculosis^7,8,9,10^, and other conditions such as Charcot Marie Tooth disease where the pathology of the disease is difficult to model in small animals^11^. In addition, bovine embryonic genome activation (EGA) has been proposed as a useful model for human EGA, supported by comparative studies^12^. To address these agricultural challenges and leverage cattle as biomedical models, it is essential to decipher the genetic and molecular architecture underlying complex phenotypes of economic, ecological, and biomedical importance in cattle.

Deciphering this architecture requires a comprehensive characterization of the regulatory genome, as it drives phenotypic variation among individuals, populations, and species. The importance of regulatory elements has been well documented in humans and model organisms^13^, where large-scale initiatives like the Encyclopedia of DNA Elements (ENCODE) and Genotype-Tissue Expression (GTEx) projects have provided fundamental insights into the molecular mechanisms of complex traits and diseases^14,15^. Paralleling these efforts, the ongoing FAANG and FarmGTEx initiatives aim to systematically annotate the regulatory landscapes across farmed animals^16,17^, such as cattle, pigs, sheep, and chickens^18–26^. Although the pilot CattleGTEx study demonstrated the utility of expression quantitative trait loci (eQTL) for dissecting complex traits in cattle^18^, its scope was constrained by limited sample size, reduced tissue coverage, and incomplete molecular phenotypes. Consequently, the majority (60-95%) of genome-wide association study (GWAS) loci could not be colocalized with detected eQTL across complex traits, giving rise to the so-called ‘missing regulation’ phenomenon similar to what has been previously observed in humans^27,28^. This limited overlap likely stems from trait-associated variants exerting regulatory effects on specific molecular phenotypes (e.g., alternative splicing and RNA stability) in specific tissues^28,29^, or possessing small effect sizes that previous datasets lack the statistical power to detect^30,31,32,33^.

To address these limitations, we report here the Phase 1 of CattleGTEx, where we have substantially expanded sample size (from 4,889 to 12,422), tissue coverage (from 23 to 43), molecular phenotype (from two to seven), and breeds (from 46 to 82) relative to the pilot study (**Fig.** 1). Our dataset achieves an average of 489 individuals per tissue (range: 52 in adrenal gland to 2,693 in immune tissues) (**Fig.** 1a-c). To enable high-resolution genetic analyses, we constructed a multi-breed genotype imputation reference panel of 3,530 individuals worldwide^34^, whose population structure closely matches that of the RNA-seq cohort (**Fig.** 1d). Using this resource, we systematically associated 4,317,531 genomic variants with seven classes of molecular phenotypes derived from RNA-seq data, followed by conditional and fine-mapping analyses to disentangle primary and non-primary regulatory effects (**Fig.** 1e). We further characterized sharing patterns of these regulatory effects across molecular phenotypes, tissues, and breeds (**Fig.** 1f). By integrating GWAS signals of 44 bovine complex traits, and signatures of selection between *Bos indicus* and *Bos taurus* as well as between dairy and beef breeds, we show the power of CattleGTEx Phase 1 for elucidating complex trait architecture, environmental adaptation, and selective breeding (**Fig.** 1h-i). Ultimately, by mapping evolutionary constraints of orthologous genes and variants between cattle and humans, we highlight the translational potential of CattleGTEx for dissecting complex traits in humans (**Fig.** 1j). All uniformly processed data, results, and computational codes are publicly available through the CattleGTEx web portal (https://cattlegtex.farmgtex.org/) (**Fig.** 1g).

**Fig 1.**
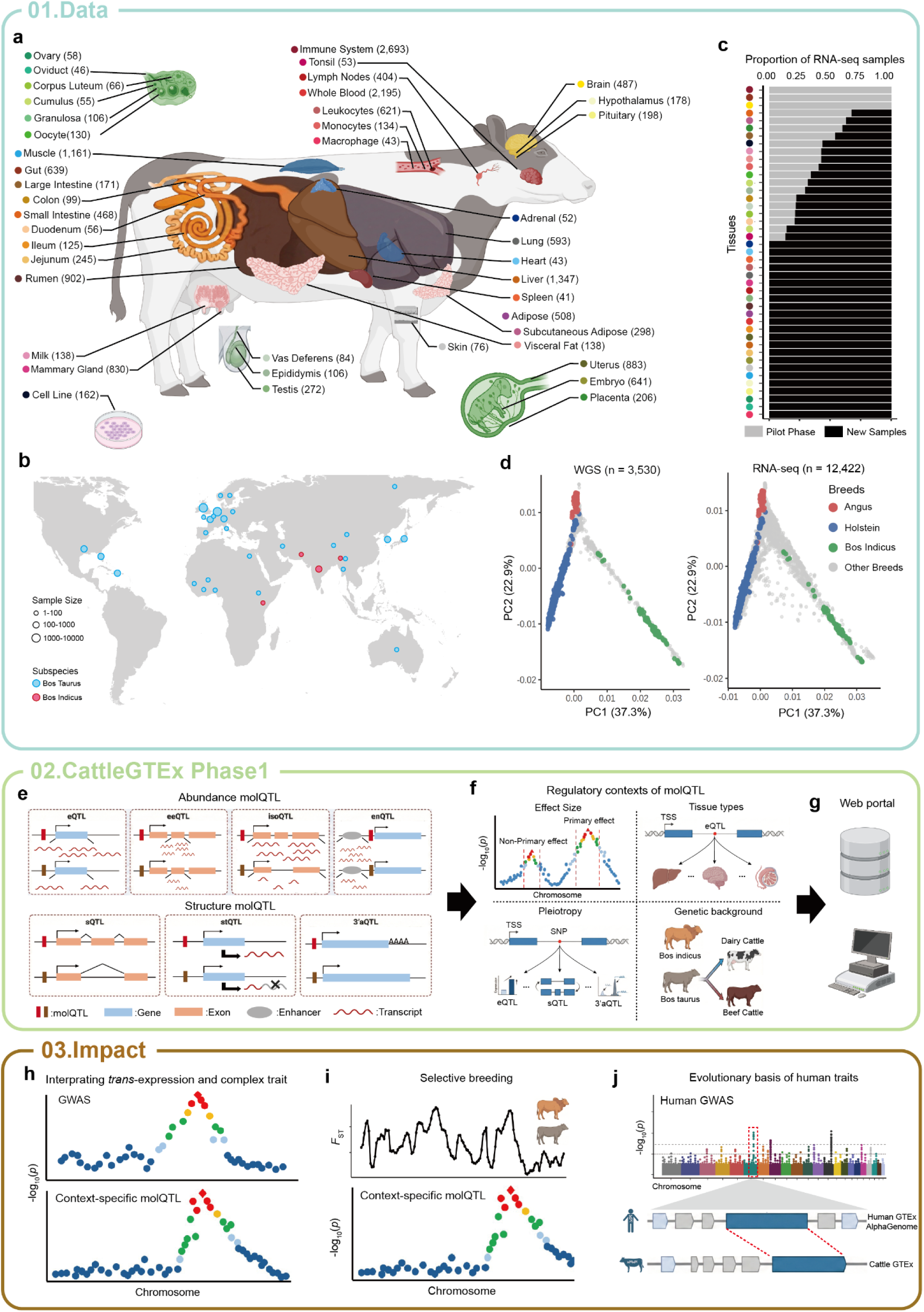
Overview of the CattleGTEx Phase1 project. **a**, Summary of 43 tissues profiled for molecular quantitative trait loci (molQTL) mapping. Sample sizes (n ≥ 40) are indicated in the respective parentheses, representing a total of 8,956 animals. The tissue colour code is used consistently throughout the manuscript. **b,** Geographic distribution of the 12,422 cattle RNA-seq samples analysed in this study. **c**, Comparison of sample sizes between the pilot study and Phase 1. Grey bars represent sample sizes in the pilot CattleGTEx project, while black bars represent newly collected samples in Phase 1. **d**, Principal component analysis (PCA) of 3,530 whole-genome sequencing (WGS) data and 12,422 RNA-seq samples. **e-g**, Overview of molQTL mapping and resource generation. We mapped seven types of molQTL and performed four types of regulatory context analyses. eQTL, expression QTL; eeQTL, exon expression QTL; isoQTL, isoform expression QTL; enQTL, enhancer expression QTL; sQTL, splicing QTL; stQTL, QTL for RNA stability of genes; 3’aQTL, QTL for 3’ UTR alternative polyadenylation (APA). All data are accessible at https://cattlegtex.farmgtex.org/. **h-j**, Applications of the CattleGTEx Phase1 resource. Analyses include interpreting genome-wide association studies (GWAS) results of 44 bovine complex traits, exploring population divergence (between *B. taurus* and *B. indicus*, as well as between dairy and beef cattle), and investigating evolutionary constraints on orthologous genes and regulatory variants between cattle and humans.

## Results

### Data summary and molecular phenotyping

In total, we collected 27,681 RNA-seq samples spanning 103 bovine tissues (**Supplementary Table** 1-2). After quality control (**Fig.** 2a**; Supplementary Fig.** 1**; Supplementary Note**), 19,789 high-quality RNA-seq samples of 88 tissues were retained for further analysis, and their genotypes were imputed using the population structure matched reference panel described above^34^. Following extensive quality control and independent validation of genotype imputation accuracy (**Supplementary Notes**), we obtained 4,317,531 common variants (minor allele frequency, MAF > 0.05), with a median concordance rate of 0.97 and genotype Pearson’s correlation-*r*² of 0.91 (**Fig.** 2b). On average, 4,576 variants were available within the *cis*-window (±1 Mb from the transcription start site, TSS) of expressed genes (Transcript per million, TPM > 0.1 and read count > 6) **(Supplementary Fig.** 2a-d). Compared with all 28,958,211 variants in the full reference panel, RNA-seq imputed variants were more enriched in gene bodies and *cis*-regulatory elements (e.g., promoter and enhancer) but depleted in intergenic regions (**Fig.** 2c**; Supplementary Fig.** 2e). After removing duplicated samples within each tissue based on identity-by-state (IBS ≥ 0.9), the final dataset comprised 12,422 RNA-seq samples from 8,956 individuals across 43 tissues, each represented by more than 40 samples **(Supplementary Fig.** 2f-g). For 5,170 samples lacking breed information, their breed identity was inferred using imputed genotypes by using a linear support vector machine (Linear SVM)^35^, achieving an overall prediction accuracy of 97.7% with 10-fold stratified cross validation **(Supplementary Fig.** 3; **Supplementary Table** 5-6).

**Fig 2.**
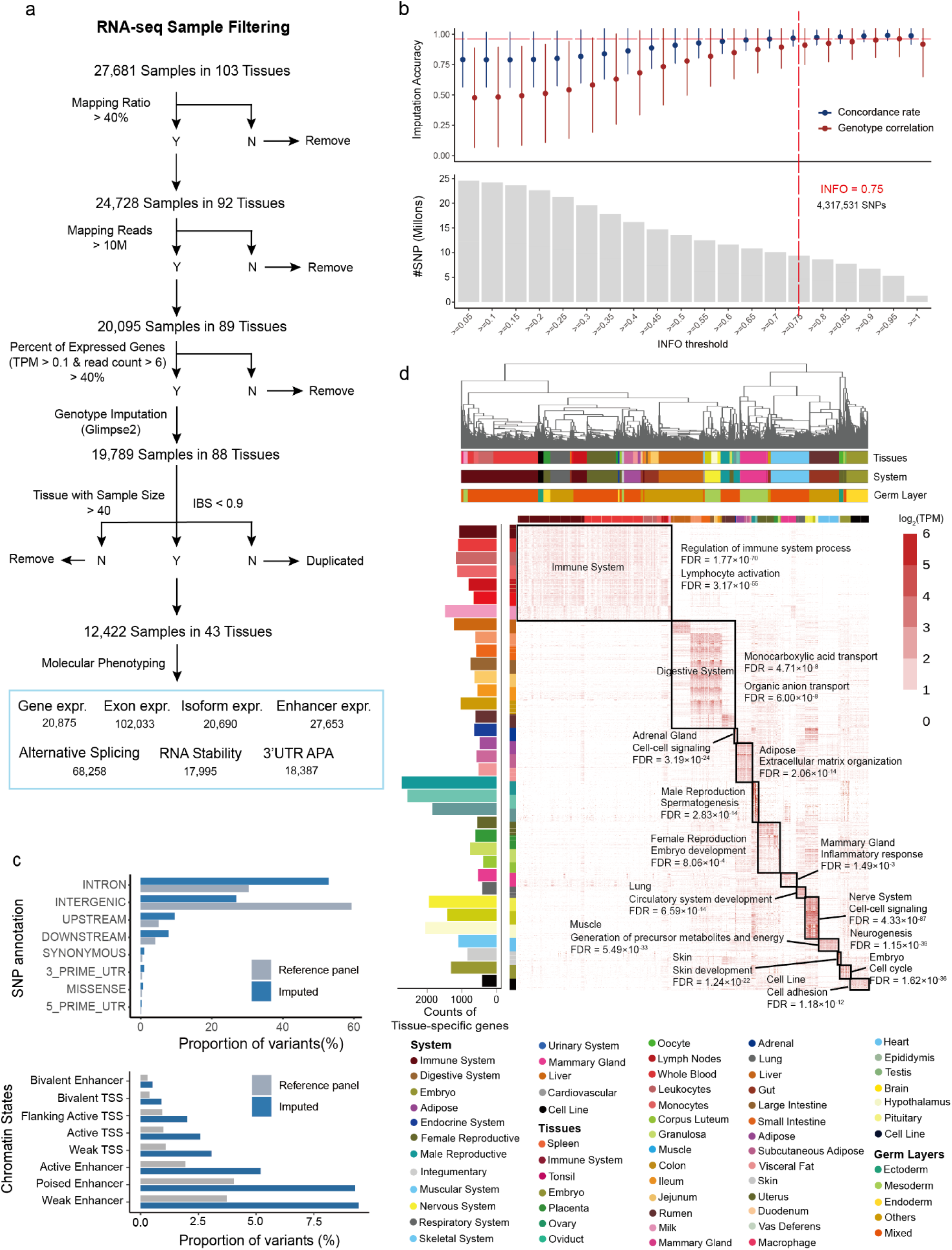
Genotype imputation and molecular phenotyping. **a**, Workflow for sample quality control (QC), duplicate removal, and molecular phenotyping. Y: Sample passed the filtering, N: Sample didn’t pass the filtering, TPM: Transcript per million, IBS: Identity-by-state, expr.: expression, APA: Alternative polyadenylation. **b**, Imputation accuracy assessed by concordance rate and genotype correlation using an external dataset, which include 15,952 samples with paired RNA-seq and whole-genome sequencing data (WGS, 30×). The top panel shows the distribution of concordance rates and genotype correlations across various INFO score cutoffs (reported by Glimpse2); the bottom panel displays the number of SNPs retained at each cutoff. **c**, Distribution of 4,317,531 imputed SNPs (from RNA-seq) and 28,958,211 reference panel SNPs across eight genomic features and eight chromatin states. TSS: Transcription start site. **d**, **Top:** Hierarchical clustering of 43 tissues based on tissue-specific gene expression, annotated by tissue type, organ system, and germ layer origin. **Bottom:** Number of tissue-specific genes (left), corresponding expression heatmap across the 43 tissues (centre), and enriched Gene Ontology (GO) terms (right).

Using these RNA-seq datasets, we defined and quantified seven classes of molecular phenotypes, including four abundance-based phenotypes (expression levels for 20,875 protein-coding and lncRNA genes, 102,033 exons, 20,690 isoforms, and 27,653 enhancers), and three structure-based phenotypes (68,258 alternative splicing, 17,995 RNA stability, and 18,387 3’UTR alternative poly-adenylation-3’UTR APA) (**Fig.** 2a). Across all molecular phenotypes, samples clustered primarily by tissue type, and then reflected developmental origin, with samples derived from the same embryonic germ layer tending to cluster together (**Fig.** 2d, **Supplementary Fig.** 4). Genes exhibiting high inter-tissue variability in expression were significantly enriched in tissue-specific biological processes, such as nervous system processes (FDR = 8.81 × 10⁻³⁰) and visual perception (FDR = 6.82 × 10⁻⁶), whereas genes with low inter-tissue variability were enriched in fundamental cellular functions such as protein localization (FDR = 3.15 × 10⁻^32^) and translation (FDR = 2.09 × 10⁻^27^) **(Supplementary Fig.** 5a, c; **Supplementary Table** 7). As expected, lncRNA genes exhibited higher tissue-specificity compared to protein-coding genes^22^ **(Supplementary Fig.** 5b). Functional enrichment analysis of tissue-specific genes (defined by FDR < 0.05 and |log₂ fold change| > 2 using the Wilcoxon rank-sum test^36^) recapitulated well-established tissue biology (**Fig.** 2d). For example, genes upregulated in immune tissues were enriched in regulation of immune system processes (FDR = 1.77 × 10⁻⁷⁰), while genes upregulated in the digestive system were enriched in monocarboxylic acid transport (FDR = 4.71 × 10⁻⁸) (**Fig.** 2d; **Supplementary Table** 8). Altogether, these results demonstrated that the derived molecular phenotypes from publicly available RNA-seq data can robustly capture tissue-specific biological programs, providing a foundation for dissecting the regulatory architecture underlying tissue-specific gene regulation.

### Fine-mapping and validation of regulatory variants

Following rigorous quality control and the mitigation of confounding factors via a linear mixed model implemented in OmiGA^37^ (**Methods**; **Supplementary Fig.** 6), we estimated the *cis*-heritability (*cis*-*h*^2^) of seven molecular phenotypes (**Supplementary Fig.** 7), and subsequently mapped molecular quantitative trait loci (molQTL) in the *cis*-windows across 43 tissues. These molQTL included four types of abundance-based signals: eQTL for gene expression, eeQTL for exon expression, enQTL for enhancer expression, isoQTL for isoform expression; and three types of structure-based signals: sQTL for alternative splicing, 3’aQTL for 3’UTR APA, and stQTL for RNA stability (**Fig.** 3a). We detected 19,180 (91.9% of tested genes) eGenes (with at least one significant variant) for gene expression, 16,358 (78.4%) eeGenes for exon expression, 7,529 (36.1%) isoGenes for isoform expression, 5,903 (28.3%) enGenes for enhancer expression, 8,454 (40.5%) sGenes for alternative splicing, 15,205 (72.9%) stGenes for RNA stability, and 7,851 (37.6%) 3’aGenes for 3’UTR APA in 35 tissues, collectively termed molGenes (**Fig.** 3a; **Supplementary Table** 9). Of them, there were 18,192 (82.9% of tested genes) genes associated with at least two types of molQTL, and 747 (3.4%) genes that were simultaneously associated with all seven molQTL. In addition, 2,203 (10%) genes were associated with a single molQTL only, including 865 eGenes, 412 eeGenes, 298 sGenes, 226 isoGenes, 183 enGenes, 147 stGenes, and 72 3’aGenes. Further statistical fine-mapping analyses for molGenes identified 771,007 credible sets (variants with cumulative PIP ≥ 95%), among which a substantial fraction contained only a single variant, such as 37.7% (85,604) of eQTL, 41.8% (75,769) of eeQTL, 29.2% (14,547) of enQTL, 35.2% (26,774) of isoQTL, 44.0% (35,710) of sQTL, 40.8% (46,077) of stQTL, and 52.3% (21,679) of 3’aQTL (**Fig**. 3b; **Extended Data Fig.** 1a-b).

**Fig 3.**
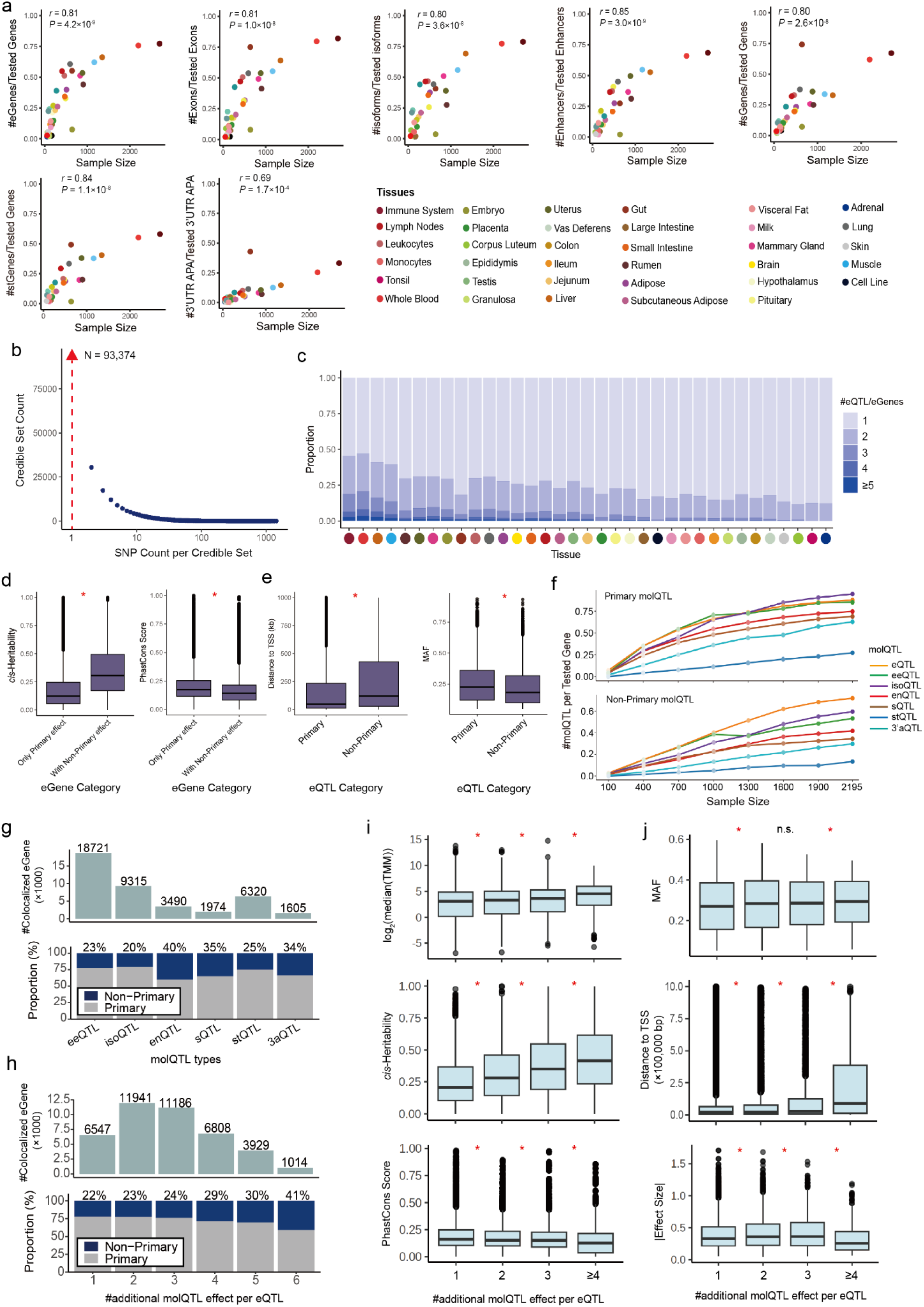
Regulatory architecture of seven molecular phenotypes across 35 tissues. **a,** Pearson’s correlation (*r*) between the proportion of molGenes (i.e., genes with significant molecular quantitative trait loci (molQTL) / all tested genes) and sample size across 35 tissues. *P* values were calculated using a two-sided Student’s *t*-test. eQTL, expression QTL; eeQTL, exon expression QTL; isoQTL, isoform expression QTL; enQTL, enhancer expression QTL; sQTL, splicing QTL; stQTL, QTL for RNA stability of genes; 3’aQTL, QTL for 3’ UTR alternative polyadenylation (APA). **b,** The size distribution of fine-mapped credible sets for all identified *cis*-eQTL across 35 tissues. The red arrow indicates 93,374 credible sets containing only one single candidate variant. **c,** The distribution of the number of conditionally independent *cis*-eQTL per gene is shown as blue stacked bars across 28 tissues. Tissues are ordered by decreasing sample size. **d,** Characteristics of genes with and without non-primary *cis*-eQTL. (Left) cis-heritability of mRNA and lncRNA expression; and (Right) sequence constraint (PhastCons score). Statistical significance was determined using a two-sided Student’s t-test. *: P < 0.05. **e,** Characteristics of primary and non-primary *cis*-eQTL. (Left) distance to the target gene; and (Right) Minor Allele Frequency (MAF). Statistical significance was determined using a two-sided Student’s t-test. *: P < 0.05. **f,** Numbers of detected primary eQTL and non-primary eQTL per tested gene across different sample size in whole blood. **g,** eQTL colocalized with other molQTL (top) and the proportion of primary versus non-primary eQTL among them (bottom). Exact numbers and proportions are shown above each bar. **h,** Pleiotropic eQTL regulating varying numbers of molecular phenotypes (top) and the proportion of primary versus non-primary eQTL among them (bottom). Exact numbers and proportions are shown above each bar. **i,** Characterization of genes harboring pleiotropic cis-eQTL. The box plots show the relationship between the number of additional molQTL regulated per lead cis-eQTL (X-axis, excluding the eQTL itself) and three properties: (Top) mRNA and lncRNA expression level, measured as log₂(median TMM) (bp); (Middle) cis-heritability of mRNA and lncRNA expression; and (Down) sequence constraint, quantified by PhastCons scores. In each box plot, the median is shown as the central line, the 25th–75th percentiles define the box limits, and whiskers extend 1.5×the interquartile range. Statistical significance was determined using a two-sided Student’s t-test. *: P < 0.05; n.s.: not significant. **j,** Characterization of pleiotropic cis-eQTL. The box plots show the relationship between the number of additional molQTL regulated per lead cis-eQTL (X-axis, excluding the eQTL itself) and three properties: (Top) Minor Allele Frequency (MAF); (Middle) distance to the target gene (bp); and (Down) absolute effect size on mRNA and lncRNA expression. In each box plot, the median is shown as the central line, the 25th–75th percentiles define the box limits, and whiskers extend 1.5×the interquartile range. Statistical significance was determined using a two-sided Student’s t-test. *: P < 0.05; n.s.: not significant.

To assess the reliability of molQTL identified above, we conducted internal and external validations, allele-specific expression (ASE) analysis at individual level, and functional enrichment (**Extended Data Fig.** 2). Internal validation across 28 tissues with sample size over 100 yielded a high replication rate (π_1_, mean = 0.93) and strong effect-size consistency (mean Pearson’s *r* = 0.89) (**Extended Data Fig**. 2a). External validation using blood eQTL in an independent population of 99 Holstein cattle achieved a decreased but remain robust replication rate of 0.76 and effect size concordance of *r* = 0.6 (**Extended Data Fig.** 2c). At the individual level, lead eQTL effect sizes were significantly correlated with ASE in whole blood (*r* = 0.66) and muscle (*r* = 0.57) (**Extended Data Fig.** 2b). Beyond statistical validation, the identified molQTL recapitulated known regulatory logic for corresponding molecular phenotypes: abundance-based signals (e.g., eQTL and isoQTL) clustered near TSS, whereas structure-based signals (e.g., sQTL and 3’aQTL) were polarized toward the transcription end site (TES) (**Extended Data Fig.** 1c-d). Notably, isoQTL exhibited the highest enrichment in start-gain and 5’UTR, while 3’aQTL and enQTL specifically targeted 3’UTR/stop-codons and enhancer-like chromatin states, respectively (**Extended Data Fig.** 1c). Finally, we verified the robustness of our approach by comparing molQTL mapping using RNA-seq imputed versus whole-genome sequencing (WGS)-called genotypes in an independent population with paired RNAseq and 30x WGS data (n = 99) (**Extended Data Fig.** 2d-m). Both approaches obtained comparable statistical power for eGene discovery and identified the same causal signals, as evidenced by lead variants being in high LD with consistent effect sizes and genomic distributions (**Extended Data Fig.** 2d-m).

### Functional and evolutionary signatures of primary and non-primary effects

To further explore the complexity of the bovine regulatory landscape beyond the lead association signals, we performed conditional analysis to identify independent variants. Across all seven molecular phenotypes, an average of 21.0–24.3% of genes harboured at least two independent molQTL (**Fig**. 3c; **Extended Data Fig.** 3). Within each molGene, we defined the variant with the strongest association as the primary effect and subsequent independent variants as non-primary effects. Notably, compared to genes with single signals, genes with multiple independent signals exhibited significantly higher *cis-h²* and lower evolutionary constraint at the sequence level (**Fig**. 3d; **Supplementary Fig.** 9), a pattern consistent with a regime of relaxed purifying selection^32^. This observation suggests that genes with more "robust" or redundant regulatory architectures might be better positioned to accumulate and maintain diverse genetic regulatory variation. Compared to primary molQTL, variants with non-primary effects were located at greater distances to the TSS and displayed lower MAF (**Fig**. 3e; **Supplementary Fig.** 9). While primary variants typically dominated core regulatory elements (e.g., 5′UTR and active promoter), non-primary molQTL showed specialized enrichments. For instance, compared to their primary effects, non-primary enQTL and 3′aQTL showed significantly higher enrichments in bivalent enhancers, while non-primary isoQTL, sQTL, and stQTL were enriched in non-coding exon variants, poised enhancer, and start-gained codon, respectively (**Extended Data Fig.** 4). These findings suggested that while primary variants govern the baseline molecular output, non-primary variants might provide a secondary layer of "fine-tuning" through distal or context-specific regulatory elements.

Finally, to evaluate the power to detect independent regulatory signals, we performed a down-sampling analyses using the whole blood samples (n = 2,195) to model the relationship between discovery power of molQTL and sample size. We observed that the detection of both primary and non-primary signals follows a non-saturated trajectory at the current sample size, with the discovery of non-primary effects showing a particularly steep dependence on sample size (**Fig**. 3f). These results highlight that a substantial portion of the bovine "fine-tuning" regulatory architecture remains hidden, and further expansion of the CattleGTEx resource will be essential to achieving a saturating map of gene expression regulation.

### Pleiotropic effects of genomic variants on molecular phenotypes

To further examine whether individual genomic variants act as multi-layer regulators or site-specific modulators (vertical pleiotropic effect), we performed colocalization analyses between eQTL and the other six molQTL. Our results revealed a fundamental decoupling between abundance-based and structure-based regulation (**Extended Data Fig.** 5a). More specifically, 39.5% of sQTL, 35.1% of 3aQTL, and 39.2% stQTL showed distinct signals (PP.H3 > 0.8) from their corresponding eQTL (**Extended Data Fig.** 5a), a finding corroborated by the presence of low LD between lead variants (LD range: 0.06–0.22) (**Extended Data Fig.** 5b). These results suggest that different layers of the transcriptomic lifecycle, such as total abundance versus isoform structure, are frequently governed by distinct genetic determinants. Examples of this decoupling include genes such as *VWA3B* in the brain and *MRPS25* in adipose tissue, where eQTL and independent structural molQTL coexist within the same tissue but operate through independent genomic variants (**Extended Data Fig.** 5d).

Despite the prevalence of layer-specific regulation, 40.0%, 20.7%, 6.2%, 5.1%, 16.6% and 10.7 % of eGenes were colocalized (PP.H4 > 0.8) with eeGenes, isoGenes, sGenes, 3aGenes, stGenes and enGenes, respectively (**Extended Data Fig.** 5c). We identified 41,425 pleiotropic eQTL (including 31,044 primary and 10,381 non-primary eQTL) that modulated at least one additional molecular phenotype, with some variants (2.4%) acting as master regulators for up to six molecular phenotypes (**Fig**. 3g). Crucially, non-primary eQTL contributed a disproportionately higher fraction of pleiotropic effects when coordinating gene expression with enhancer activity (enQTL) or structural phenotypes, whereas primary eQTL dominated the regulation of abundance-based phenotypes such as exon and isoform expression (**Fig**. 3g). Furthermore, the contribution of non-primary eQTL increased progressively with the degree of pleiotropy (**Fig**. 3h). This suggest that while primary eQTL drive the majority of total expression variance, non-primary eQTL likely play a key role in coordinating the interplay between expression and post-transcriptional modulation (such as splicing and RNA stability). Genes regulated by genomic variants with higher pleiotropic effects were characterized by elevated expression levels, higher *cis-h²*, and reduced evolutionary constraint, mirroring the "relaxed selection" profile observed for complex regulatory architectures (**Fig**. 3i). The respective pleiotropic variants displayed higher MAF, larger effect sizes on gene expression, and stronger enrichment in regulatory regions (e.g., 5′UTRs, 3′UTRs, promoters, and enhancers) (**Fig**. 3j; **Extended Data Fig.** 5e).

### Tissue-sharing patterns of regulatory effects

In general, tissues with similar biological functions (e.g., within the nervous or digestive systems) tended to cluster together based on their molQTL effect profiles (**Fig.** 4a; **Supplementary Fig**. 10). However, rumen, a ruminant-specific digestive organ, clustered separately from other intestinal regions across all molQTL types, underscoring their specialized cellular composition, developmental origin and biological roles in plant-based digestion^38^. These regulatory effects tended to be either tissue-specific or ubiquitous across tissues, consistent with previous findings in other species^18–20,22^ (**Fig**. 4b). The eQTL that were active in more tissues (stronger horizontal pleiotropic effects) exhibited smaller effect sizes, higher MAF, shorter distances to TSS, and more enrichment in regulatory elements (e.g., 3’UTR and 5’UTR) (**Supplementary Fig**. 11). Notably, primary regulatory effects showed a slightly greater tissue specificity than non-primary ones (**Fig**. 4c; **Supplementary Fig**. 12a). Furthermore, eQTL with stronger vertical pleiotropic effects tended to be more tissue-specific (weaker horizontal pleiotropic effects) (**Fig**. 4d; **Supplementary Fig**. 12b), indicating that complex regulatory mechanisms are often confined to specific tissue contexts.

**Fig. 4.**
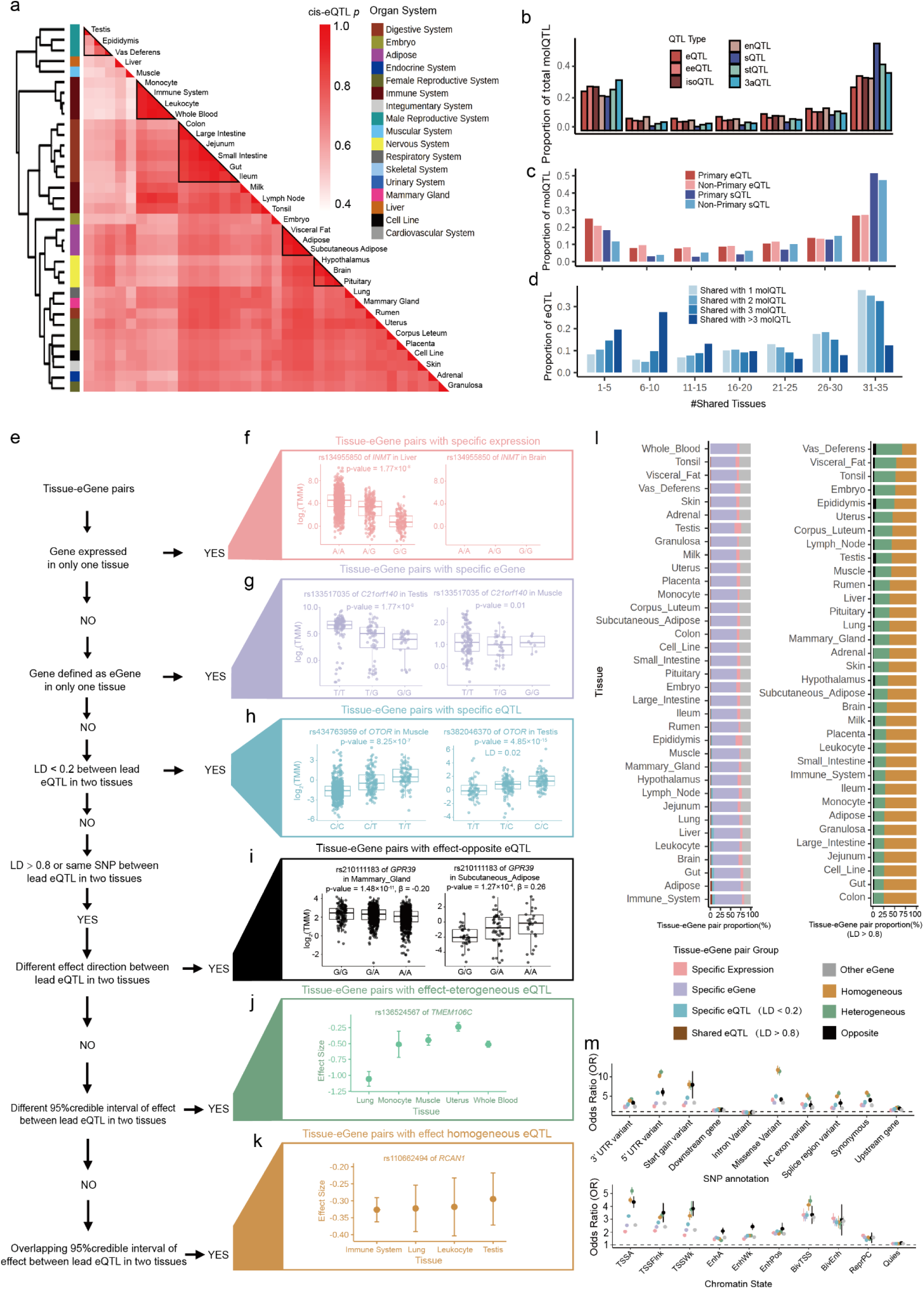
Tissue-specific regulatory effects. **a,** Heatmap showing pairwise Spearman’s correlation (*ρ*) of *cis*-eQTL effect sizes across 51 tissues. Tissues were hierarchically clustered using the complete linkage method based on the maximum distance of *ρ.* **b,** Proportion of molQTL active across tissues, measured by the number of tissues with a *mashr* local false sign rate (LFSR, equivalent to FDR) < 0.05. **c,** Proportion of primary and non-primary *cis*-eQTL and *cis*-sQTL active across different numbers of tissues. **d,** Proportion of *cis*-eQTL shared with different numbers of other molQTL active across different numbers of tissues. **e,** Pipeline for classifying different tissue-eGene pairs in each tissue. LD means linkage disequilibrium. **f,** Example of expression-specific tissue-eGene pairs, including gene name, lead eQTL, p-value and three genotypes in gene-expressed tissue (left) and non-expressed tissue (right). The two-sided P value of SNPs was calculated by the linear mixed regression model. **g,** Example of eGene-specific tissue-eGene pairs, including gene name, lead eQTL, p-value and three genotypes in eGene-defined tissue (left) and non-eGene tissue (right). The two-sided P value of SNPs was calculated by the linear mixed regression model. **h,** Example of eQTL-specific tissue-eGene pairs, including gene name, lead eQTL, p-value, three genotypes and LD between lead eQTL from two tissues. The two-sided P value of SNPs was calculated by the linear mixed regression model. **i,** Example of effect-opposite tissue-eGene pairs, including gene name, lead eQTL, p-value, three genotypes and effect sizes from lead eQTL in two tissues. The two-sided P value of SNPs was calculated by the linear mixed regression model. **j,** Example of effect-heterogeneous tissue-eGene pairs. The title represents gene name and lead eQTL. Y axis represents the 95% credible interval of effect size in lead eQTL. The first tissue in X axis represents the tested tissue and others represent tissues with non-overlapped interval with the tested tissue. **k,** Example of effect-homogeneous tissue-eGene pairs, The title represents gene name and lead eQTL. Y axis represents the 95% credible interval of effect size in lead eQTL. The first tissue in X axis represents the tested tissue and others represent tissues with overlapped interval with the tested tissue. **l,** Proportion of each tissue-eGene classification. Bar plot in the left represents proportion of expression-specific, eGene-specific, eQTL-specific (LD < 0.2), eQTL-shared (LD > 0.8) and other tissue-eGene pairs, whereas the right one represents proportion of eQTL-shared (LD > 0.8) tissue-eGene pairs, including effect-opposite, effect-heterogeneous and effect-homogeneous. **m,** Functional enrichment between each group of tissue-eGene and regulatory elements (top: SNP classification, bottom: chromatin states).

To systematically characterize tissue specificity of eQTL between any two tissues, we first identified three classes of tissue-specific eGenes (**Fig**. 4e) : **1) Specific-expression**, defined as genes expressed in only one of each of the tissue pairs (e.g., *INMT* between liver and brain, **Fig**. 4f); **2) Specific-eGenes**, defined as detected eGenes in only one tissue despite being expressed in both tissues (e.g., *C21of140* between testis and muscle, **Fig**. 4g); **3) Specific-eQTL**, defined as lead variants of shared eGenes were weakly linked (LD < 0.2), such as *OTOR* between muscle and testis (LD = 0.02; **Fig**. 4h). We defined shared-eQTL when lead variants of same eGenes were highly correlated (LD ≥ 0.8) (**Fig**. 4e). These shared-eQTL were further divided into three classes according to their direction and magnitude of effect sizes between tissues: **1) Opposite**, eQTL exhibiting opposite effect directions between tissues (e.g., *GPR39* between mammary gland and adipose, **Fig**. 4i); **2) Heterogeneous**, eQTL with concordant effect directions but distinct effect size (non-overlapping 95% confidence intervals for effect size estimates), such as *TMEM106C* between lung and other tissues (**Fig**. 4j); **3) Homogeneous**, eQTL with overlapping confidence intervals (e.g., *RCAN1* across immune system tissue and lung, leukocyte, and testis (**Fig**. 4k).

Among all eGenes, **Specific-eGenes** constituted the largest proportion (64%) across tissues, followed by **Specific-expression** (7%), and **Specific-eQTL** (2%, **Fig**. 4l). Among shared-eQTL, **Homogeneous** constituted the largest proportion across tissues (63%), followed by **Heterogeneous** (33%) and **Opposite** (4%). These shared-eQTL showed shorter distances to TSS, higher MAF, larger effect sizes, and stronger enrichment in active regulatory elements compared with other eQTL (**Fig**. 4m; **Supplementary Fig**. 13). Notably, among shared-eQTL, **opposite** showed a strongest enrichment in enhancers and flanking TTS regions (**Fig**. 4m), indicating that these variants may participate in context-dependent or antagonistic regulatory mechanisms that fine-tune gene expression in a tissue-specific manner.

Given the widespread tissue-sharing of eQTL, particularly those with small effect size, we conducted a joint *cis*-eQTL mapping analysis by pooling 8,743 non-redundant samples across all tissues (see **Supplementary Notes**). This integrative approach identified 30,880 additional fine-mapped eQTL (variants with cumulative PIP ≥ 95%) that were previously undetected in single-tissue analyses. Compared to eQTL detected via the single-tissue framework, these newly discovered variants were characterized by smaller effect sizes, lower MAF, greater distances to TSS, and higher enrichment in enhancers (**Supplementary Fig**. 14).

### Deciphering the regulatory architecture underlying population selection signatures

To explore the regulatory conservation between *B. taurus* and *B. indicus*, we conducted population-stratified eQTL mapping across eight tissues, each with at least 40 individuals for both two subspecies (**Methods**; **Supplementary Fig**. 15a-b). In general, effect sizes of population-stratified eQTL clustered primarily by tissue types and then by subspecies (**Fig**. 5a). The eQTL showed higher replication rates (mean π₁ = 0.73) and effect-size concordance (mean Spearman’s *ρ* = 0.76) between subspecies than between tissues (**Fig**. 5a; **Extended Data Fig.** 6a). Even though they diverged at least 300,000 years ago, these results highlighted the conservation of regulatory effects between the two subspecies^39–41^.

**Fig. 5.**
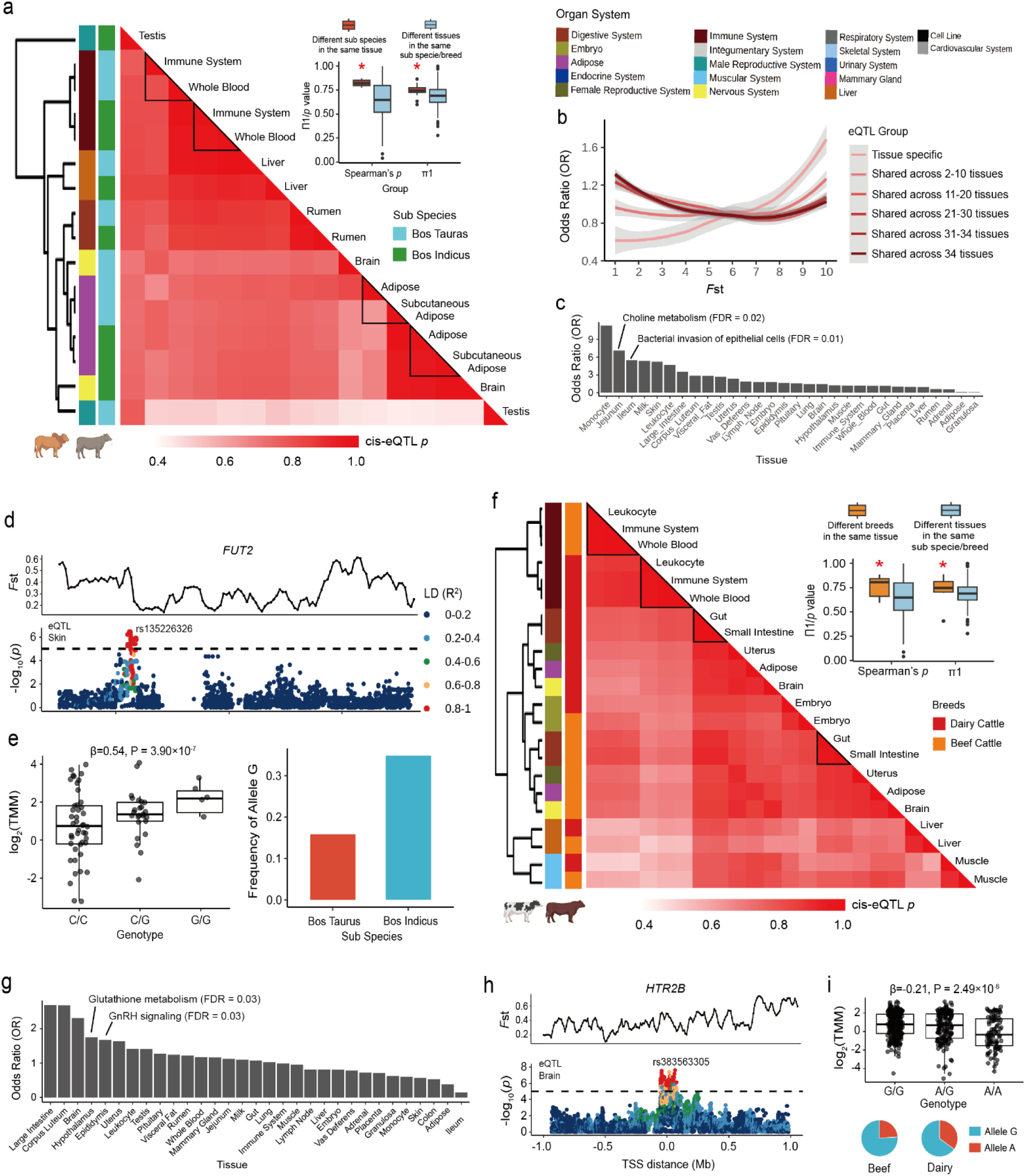
Liking molQTL to selection signatures between subspecies and breeds. **a,** Heatmap showing pairwise Spearman’s correlation (*ρ*) of population-stratified *cis*-eQTL effect sizes across 8 tissues between *B. taurus* and *B. indicus*. Tissues were hierarchically clustered using the complete linkage method based on the maximum distance of *ρ.* Statistical significance of Π1 and p-value between two groups were determined using a two-sided Student’s t-test. *: P < 0.05. **b,** Enrichment (odds ratio) of shared and tissue-specific molQTL across 10 *F*_ST_ deciles. The lines were fitted by using the geom_smooth function from ggplot2 (v3.3.2) in R (v4.0.2). **c,** Enrichment (odds ratio) of tissue-specific eQTL and strongly selected regions (top 5% *F*_ST_) between *B. taurus* and *B. indicus* across 35 tissues. Associated KEGG pathways related to immune response are indicated. **d,** Manhattan plots illustrating an example of a selected eGene (*FUT2*) in skin. The eQTL rs135226326 lies in a high-*F*_ST_ region (*F*_ST_ = 0.54). The two-sided P value of lead SNP was calculated by the linear mixed regression model. **e,** Functional interpretation of rs135226326: boxplot (left) shows expression of *FUT2* across three genotypes (C/C, C/G and G/G), while the barplot (right) shows the frequency distribution of the allele G between *B. taurus* and *B. indicus*. The two-sided P value of lead SNP was calculated by the linear mixed regression model. **f,** Heatmap showing pairwise Spearman’s correlation (*ρ*) of population-stratified *cis*-eQTL effect sizes across 11 tissues between dairy and beef cattle. Tissues were hierarchically clustered using the complete linkage method based on the maximum distance of *ρ.* Statistical significances of Π1 and p-value between two groups were determined using a two-sided Student’s t-test. *: P < 0.05. **g,** Enrichment (odds ratio) of tissue-specific eQTL and strongly selected regions (top 5% *F*_ST_) between dairy and beef cattle across 35 tissues. Associated KEGG pathways related to growth and development are indicated. **h,** Manhattan plots illustrating an example of a selected eGene (*HTR2B*) in brain. The eQTL rs383563305 lies in a high-*F*_ST_ region (*F*_ST_ = 0.58). The two-sided P value of lead SNP was calculated by the linear mixed regression model. **i,** Functional interpretation of rs383563305: boxplot (top) shows expression of *HTR2B* across three genotypes (G/G, G/A and A/A), while the pie chart (bottom) shows the frequency distribution of the allele G and allele A between dairy and beef cattle. The two-sided P value of lead SNP was calculated by the linear mixed regression model.

To further explore the regulatory adaptation of *B. indicus* to heat, humidity and parasites challenges in tropical environments, we scanned the genome-wide selection sweeps between 200 *B. taurus* and 142 *B. indicus* with WGS data (> 10×) via an *F*_ST_-based approach^34^ (**Supplementary Table** 11; **Methods**). The 248,679 genomic regions tested were evenly divided into 10 deciles (*F*_ST_1-10) based on their *F*_ST_ values from the smallest (no selection) to the largest (strongest selection). For example, the *F*_ST_10 genomic region contained *DNAJC18* (*F*_ST_ = 0.86), *HOXC4* (*F*_ST_ = 0.77), and *FAAP20* (*F*_ST_ = 0.77), which were previously reported to be implicated in environmental adaptation between cattle populations^42–44^ (**Supplementary Fig**. 16a). The enrichment of fine-mapped molQTL in each of 10 deciles was then tested. Notably, tissue-specific molQTL showed a significantly higher enrichment in regions with larger *F*_ST_ values (**Fig**. 5b; **Supplementary Fig**. 16b). As expected, tissues with higher enrichment of eQTL in strongly selected regions showed lower correlation of effect-size between *B. taurus* and *B. indicus* in the population-stratified analysis (**Extended Data Fig.** 6b). Among all tissues, molQTL specific in skin, adipose, immune and digestive tissues displayed the highest enrichment in the *F*_ST_10 group (**Fig**. 5c; **Supplementary Fig**. 16c). The eGenes of these tissues located in strongly selected regions were significantly involved in immune-related functions such as choline metabolism (FDR = 0.02) and bacterial invasion (FDR = 0.01; **Fig**. 5c). For example, the lead eQTL (rs135226326) of *FUT2*, a skin-specific eGene, resided in a strong selection region (*F*_ST_ = 0.54; **Fig.** 5d). The G allele of rs135226326 was associated with increased *FUT2* expression and occurred at a higher frequency in *B. indicus*, potentially enhancing immune responsiveness in extreme environments and thereby contributing to adaptive differentiation between *B. taurus* and *B. indicus* due to the role of *FUT2* in mediating antiviral immune defence^45^. These results suggest that regulatory variants in immune and metabolic tissues have played important roles in the divergence of *B. taurus* and *B. indicus*, likely reflecting selection pressures related to environmental, metabolism, and immune defence adaption^2^.

To elucidate the regulatory mechanisms underlying intensive artificial selection between dairy and beef breeds in more recent timescales, we conducted population-stratified eQTL in 11 tissues (sample size > 40 in each of breed**; Supplementary Fig.** 15c-d). Similar to findings between *B. taurus* and *B. indicus*, effect sizes of eQTL were generally conserved between dairy and beef (**Fig**. 5f; **Extended Data Fig.** 7). Compared to tissue-shared eQTL, tissue-specific ones also showed significantly higher enrichment in strongly selected regions between 651 dairy and 552 beef cattle detected using WGS data (**Supplementary Fig**. 17a; **Supplementary Table** 12). In contrast to the divergence between *B. taurus* and *B. indicus*, tissues with highest enrichment of eQTL in strongly selected regions between dairy and beef were brain-related tissues, particularly hypothalamus, rather than skin and immune tissues (**Fig**. 5g; **Supplementary Fig**. 17b). Brain-specific eGenes located in the strongly selected regions (*F*_ST_10) were significantly enriched in pathways related to growth and developmental regulation, including glutathione metabolism (FDR = 0.03) and the GnRH signalling pathway (FDR = 0.03; **Fig**. 5g). This indicated that artificial selection between dairy and beef cattle has preferentially targeted neuroendocrine and growth-regulatory mechanisms. For example, the lead eQTL rs383563305 for *HTR2B*, a brain-specific eGene, resided in a region under strong selection (*F*_ST_ = 0.58; **Fig**. 5h). The allele G in rs383563305 was associated with upregulated expression of *HTR2B* and showed a higher frequency in beef compared to dairy cattle (**Fig**. 5i). As *HTR2B* mediates serotonergic regulation of hypothalamic appetite control, selection on its eQTL likely reflects divergent breeding pressures on feed intake, energy allocation and growth-related traits between beef and dairy cattle^46^. Altogether, these results suggest that both long-term adaptive evolution and intensive artificial breeding tended to target on variants with tissue-specific effects rather than those with ubiquitous effects.

### The genetic and molecular architecture underlying bovine complex traits

To investigate the regulatory mechanisms underlying complex traits in cattle, we examined 2,070 GWAS loci of 44 complex traits of economic importance, encompassing 22 body conformation, 6 milk production, 8 reproduction, and 8 health traits (**Supplementary** Table 13). We systematically explored enrichment patterns of these GWAS loci against all seven types of molQTL and different categories of regulatory variants based on effect size, tissue-sharing pattern, and pleiotropy. Overall, eQTL showed the strongest enrichment with GWAS loci, followed by eeQTL, 3’aQTL, sQTL, isoQTL, and stQTL (**Fig**. 6a). Notably, compared to primary effects, non-primary effects exhibited higher enrichment in GWAS loci, consistent across all types of molQTL (**Fig**. 6a). Compared to tissue-shared effects, tissue-specific effects were more enriched in GWAS loci (**Fig**. 6a; **Extended Data Fig.** 8a). The eQTL with stronger pleiotropy showed higher enrichment in GWAS loci (**Fig**. 6a). These observations were further supported by their similarity to GWAS loci in terms of distance to TSS, population selection pressure (measured by MAF-effect size relationship), and enrichment in regulatory elements (**Fig**. 6b-d; **Extended Data Fig.** 8b-c). We thereby propose a genetic regulatory model in which genetic effects spanning multiple layers of the central dogma (vertical pleiotropic effects) are more likely to propagate to complex traits, whereas variants acting across multiple biological contexts of a single biological layer (horizontal pleiotropic effects) may incur greater fitness costs and are therefore less likely to translate their effects to higher-order phenotypes (**Fig**. 6e).

**Fig. 6.**
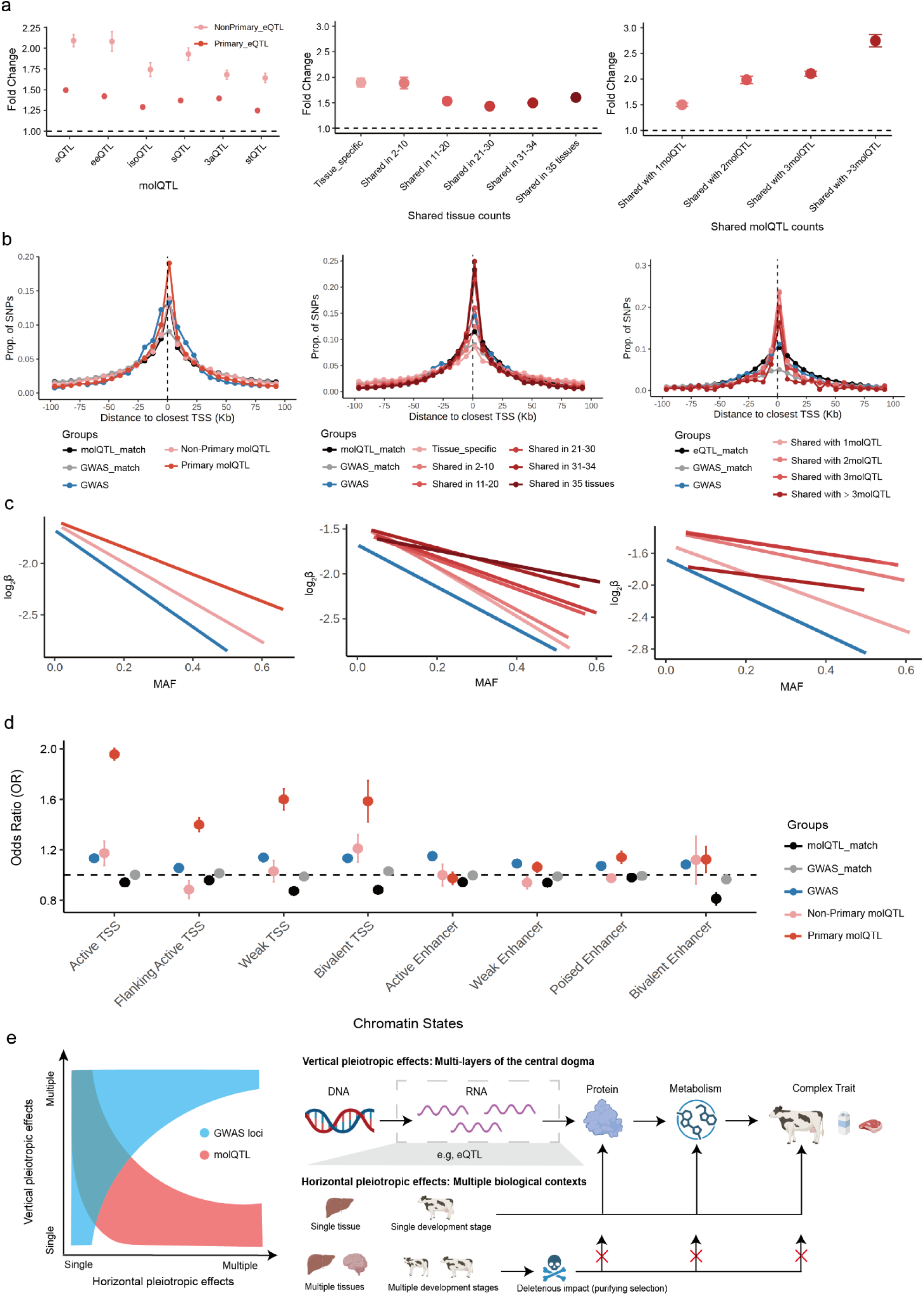
Enrichment between three classes of molQTL and GWAS loci from 44 complex traits. **a,** Enrichment (odds ratio) of three classes of molQTL with genome-wide associations (GWAS) of 44 cattle traits using QTLenrich (v.2.0), including primary and non-primary molQTL (**left**), tissue-sharing eQTL (**mid**) and pleiotropic eQTL (**right**). **b**, Distance to nearest TSS regions of three classes of molQTL, GWAS loci, MAF-matched GWAS loci and MAF-matched molQTL, including primary and non-primary molQTL (**left**), tissue-sharing molQTL (**mid**) and pleiotropic eQTL (**right**). **c,** Selection pressure (measured by MAF-effect size relationship) of three classes of molQTL and GWAS loci, including primary and non-primary molQTL (**left**), tissue-sharing molQTL (**mid**) and pleiotropic eQTL (**right**). **d,** Functional enrichment between primary/non-priamry molQTL and regulatory elements. **e,** A genetic regulatory model of complex traits. The left plot represents the distribution of molQTL (red) and GWAS loci (blue) across different levels of biological contexts (x-axis) and layers of the central dogma (y-axis). The right plot represents the process of how regulatory genetic variants under multiple layers of the central dogma and specific biological contexts shapes GWAS signals.

To pinpoint putative causal variants and genes governing complex traits, we performed colocalization analyses of molQTL and GWAS loci. Overall, 75% (1,545) of GWAS loci, ranging from 33% (5 loci) for calving-to-conception interval to 96% (96 loci) for milk protein percentage, were colocalized (PP4 > 0.8) with at least one molQTL (**Supplementary Fig.** 18a). Each molQTL type also made unique contributions to GWAS colocalization. Among them, eeQTL accounted for the largest number of unique colocalizations (n = 64), followed by eQTL (29), stQTL (24), sQTL (21), isoQTL (17), enQTL (15), and 3′aQTL (12; **Supplementary Fig.** 18b). For instance, an isoQTL of *PHACTR2* in whole blood was exclusively associated with the udder cleft trait, while an sQTL of *NUDT6* in the corpus luteum was exclusively associated with rump width trait (**Extended Data Fig.** 9). Together, these findings underscored the importance of integrating multiple molecular phenotypes to elucidating the molecular basis of complex traits. Consistent with enrichment analyses, non-primary, tissue-specific, and pleiotropic eQTL showed higher colocalization proportions (PP4 > 0.8) with GWAS loci compared to their counterparts (**Fig**. 7a). Each category also made distinct contributions to explaining GWAS signals: non-primary eQTL contributed 57 additional colocalizations beyond primary eQTL, tissue-specific eQTL contributed 79 unique colocalizations, and pleiotropic eQTL accounted for 34 unique colocalizations (**Fig**. 7b). For example, a colocalization (PP4 = 0.93) between rump width and *EPN3* in the liver was detected for a non-primary eQTL (rs378415691) after conditioning on the primary eQTL (rs379113614). As an endocytic adaptor protein, *EPN3* regulates lipid transport and signal transduction, potentially influencing growth and development in cattle^47^. For pleiotropic eQTL, *GIGYF1*, which is related to adverse metabolic health related to higher glycemic levels and fat mass^48^, showed high colocalization with the net merit trait through both eQTL (PP4 = 0.96) and stQTL (PP4 = 0.93) effects driven by the same lead variant (rs109407080). In addition, the lead eQTL rs135173214 of *ENSBTAG00000050359* exhibited strong colocalization with somatic cell score (SCS) specifically in the mammary gland (PP4 = 0.87) but not in other tissues (e.g., adipose PP4 = 2.4×10^-2^), highlighting tissue-specific regulatory control of this gene in mastitis-related traits (**Fig**. 7c).

**Fig. 7.**
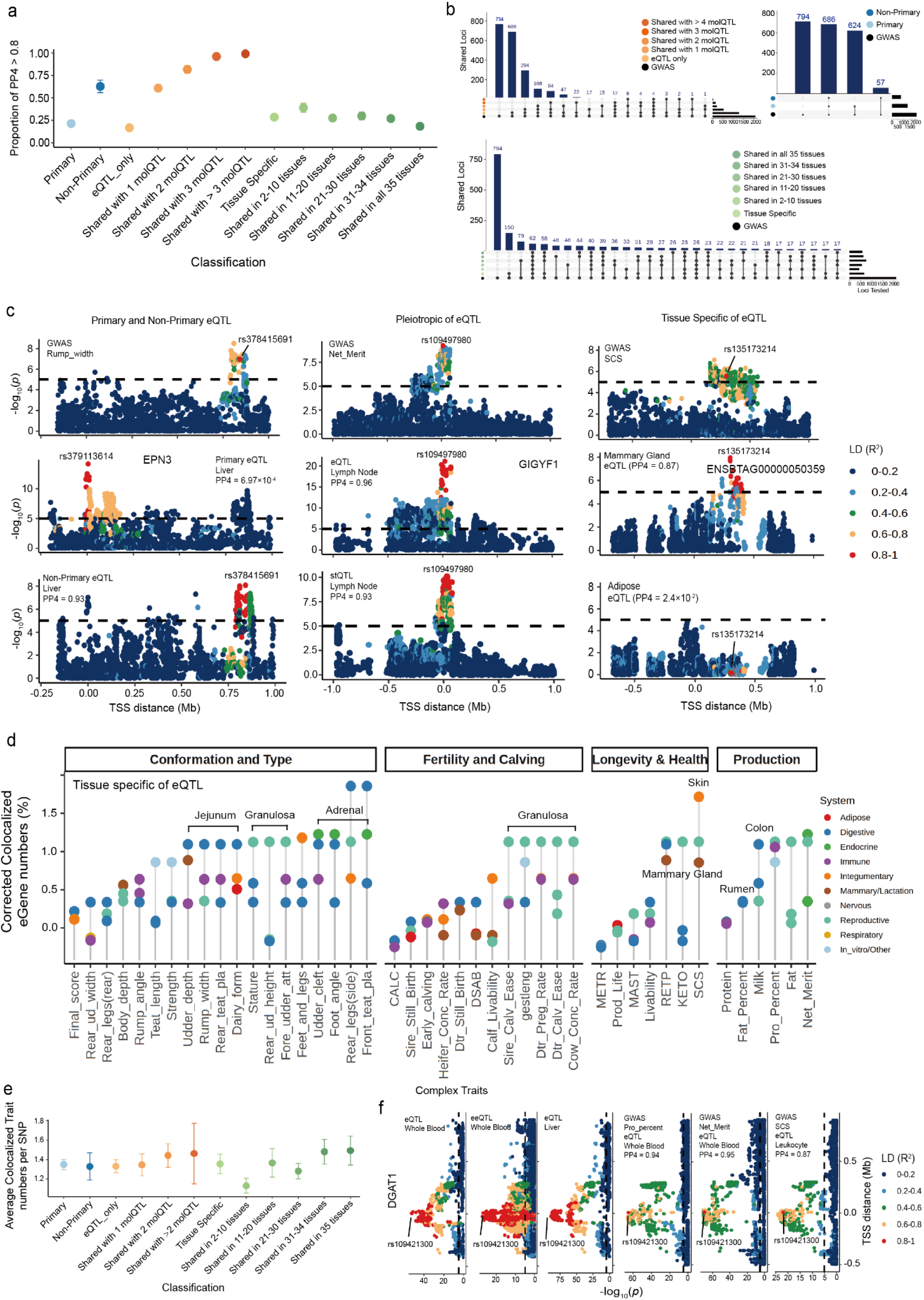
Colocalization between three classes of molQTL and GWAS loci from 44 complex traits. **a,** Proportion of colocalized molQTL (PP.H4 > 0.8) in three classes molQTL, including primary and non-primary molQT, tissue-shared molQTL and pleiotropic eQTL. **b**, Numbers of colocalized GWAS loci by each class of molQTL, including primary and non-primary molQTL (**right and top**), tissue-shared molQTL (**bottom**) and pleiotropic eQTL (**left and top**). Each blue bar represents a combination of different molQTL in each class. Each black bar (right) represents the number of GWAS loci colocalized in the respective combination. **c,** Manhattan plots showing the -log₁₀(p) distribution of specific associated signals between GWAS loci and three classes of molQTL, including rump width (GWAS) and non-primary eQTL in liver across genetic variants within the locus of *EPN3*, net merit (GWAS) and pleiotropic eQTL (with stQTL) in lymph nodes across genetic variants within the locus of *GIGYF1*, and SCS (GWAS) and tissue-specific eQTL in mammary gland across genetic variants within the locus of *ENSBTAG00000050359*. Variants are colored according to their linkage disequilibrium (LD, r²) with the lead SNP. The two-sided P values of SNPs were calculated by the linear mixed regression model. **d**, Corrected colocalized eGene counts (PP.H4 > 0.8) between each tissue and each complex trait in eQTL results. Colocalized eGene counts were corrected by detected eQTL counts. **e,** Average colocalized trait counts per SNP (PP.H4 > 0.8) in three classes of molQTL, including primary and non-primary molQT, tissue-shared molQTL and pleiotropic eQTL **f,** Manhattan plots showing the -log₁₀(p) distribution of the association signal between pleiotropic GWAS loci and molQTL for protein percent, SCS and net merit (GWAS) with eQTL and eeQTL in whole blood and liver across genetic variants within the locus of *DGAT1*. Variants are colored according to their linkage disequilibrium (LD, r²) with the lead SNP rs109421300. The two-sided P values of lead SNPs were calculated by the linear mixed regression model.

To further prioritize tissues relevance to complex traits, we adjusted the number of colocalized genes by the total number of molGenes detected in each tissue. Granulosa cells, which play a central role in follicular growth and hormonal signalling^49^, showed high relevance with multiple complex traits, including body conformation traits (e.g., stature and rear udder attachment) and reproductive traits (e.g., sire calving ease and gestation length). Mammary gland and skin exhibited high relevance with milk production and health traits, such as livability and SCS. Colon and rumen showed high relevance with milk production traits, including milk yield and fat percentage (**Fig**. 7d; **Supplementary Fig.** 19). These findings were generally consistent across all types of molQTL (**Supplementary Fig.** 18c).

To explore the molecular architecture underlying pleiotropic effects of GWAS loci, we detected 151 loci associated with at least two complex traits (**Supplementary Table**. 14). Overall, eQTL with stronger pleiotropic effects and broader tissue-sharing patterns showed higher colocalization rate with GWAS loci associated with more complex traits (**Fig**. 7e). For example, rs109421300 was associated with the gene and exon expression of *DGAT1* in whole blood and liver, a gene with major effect on milk yield and composition in cattle^50^, and showed strong colocalization with multiple complex traits, including protein percentage, net merit, and SCS (**Fig**. 7f). In addition, the lead SNP (rs110878911) that was associated with both gene and isoform expression of *LSP1* in leukocytes and liver, showed strong colocalization with multiple complex traits, including calving ease and body strength (**Supplementary Fig.** 20). *LSP1* encodes a protein that regulates immune cell activity and is involved in lymphocyte migration, activation, and intercellular interactions^51^. This gene has also been linked to the development of multiple diseases in humans, including rheumatoid arthritis, Crohn’s disease, and various cancers^52^.

### Evolutionary constraint of regulatory effects between cattle and humans

To explore the evolutionary constraint of regulatory effects, we aligned 198,897 fine-mapped cattle eQTL to the human genome *via* LiftOver (referred to as orthologous variants) and then systematically evaluated the impact of these orthologous variants on genome function and complex traits in humans (**Fig.** 8a; See **Methods)**. Compared to MAF-matched random variants (51.27%) in cattle, fine-mapped eQTL exhibited a higher mapping rate to the human genome, with 53.25∼74.45% across different groups of eQTL, indicating cross-species conservation of regulatory variants at the DNA sequence level. Primary eQTL (57.53%) were more evolutionarily constrained than non-primary ones (54.34%). Notably, variants with higher pleiotropic effect were more evolutionarily constrained (**Fig.** 8b). We then used *AlphaGenome* to explore the functional impact of these orthologous variants in the human genome^53^. Compared with MAF-matched random variants, orthologous variants of cattle eQTL exhibited significantly higher predicted quantile scores, indicative of the functional conservation of regulatory variants between cattle and humans. Orthologous variants of tissue-shared eQTL showed higher *AlphaGenome*-predicted effects than those of tissue-specific variants, and variants with higher pleiotropic effect on molecular phenotypes also displayed higher *AlphaGenome*-predicted effects (**Fig.** 8c). To test whether orthologous variants of cattle eQTL tended to overlap the same human regulatory elements as eQTL detected in human GTEx^54^, we assessed their enrichment across 15 human chromatin states predicted from ENCODE data^55^. Notably, human-orthologous variants of cattle eQTL exhibited an enrichment profile across these chromatin states that closely resembled that of human GTEx eQTL (Pearson’s *r* = 0.809, *P*-value = 2.62e-4) (**Fig. 8d; Supplementary Fig.** 22a-d). Notably, variants with higher pleiotropic effect exhibited a higher enrichment in CTCF and enhancers (**Supplementary Fig.** 21d).

**Fig 8.**
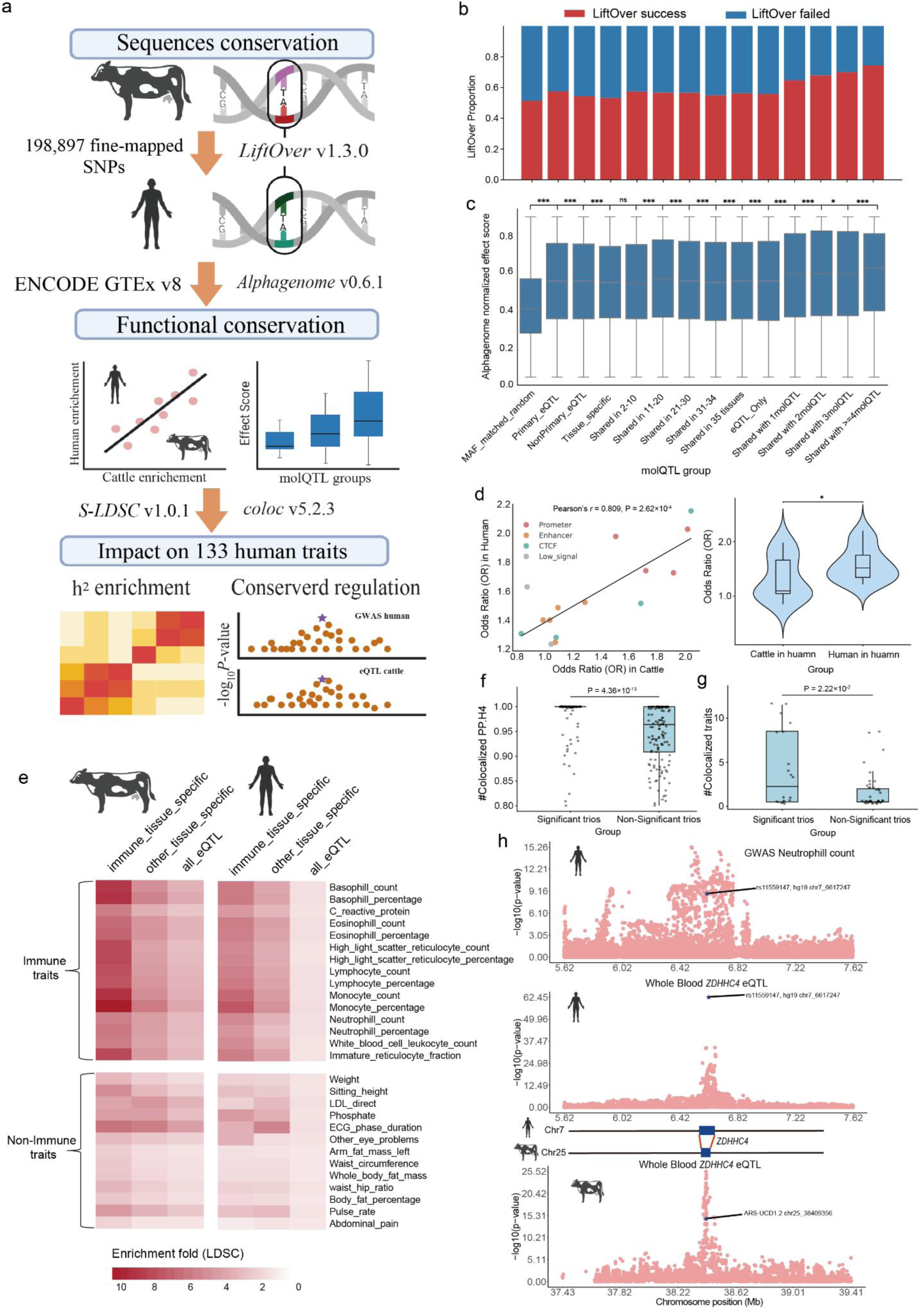
Cross-species analysis identifies conserved regulatory effects between cattle and humans. **a,** Overview of the cross-species analysis workflow. At the sequence-conservation level, fine-mapped cattle expression quantitative trait loci (eQTL) were projected onto the human genome using LiftOver. Functional conservation was assessed using ENCODE regulatory annotations and effect-score predictions from the AlphaGenome model. Conserved regulatory activity and its contribution to human complex traits were evaluated using 133 genome-wide association study (GWAS) summary statistics from UK Biobank and human GTEx expression data. **b,** Stacked bar plot showing the proportion of cattle eQTL from different groups, and a minor-allele-frequency (MAF)-matched random set, that could be successfully lifted over to the human genome. **c,** Box plots of standardized AlphaGenome-predicted effect scores for lifted-over variants from different cattle QTL groups; only tissue-level, gene-expression–related tracks were included. P values were calculated using two-sided unpaired t tests. **d,** Left, correlation between enrichment of lifted-over cattle eQTL in human ENCODE regulatory elements and enrichment of human GTEx eQTL in the same elements. Right, violin plots comparing the magnitude and distribution of enrichment estimates between lifted-over cattle eQTL and human GTEx eQTL. **e,** Stratified LD score regression (S-LDSC) heritability enrichment of tissue-specific eQTL annotations from cattle GTEx and human GTEx across complex traits; the heat map shows enrichment fold change. **f,** For 344 cattle–human orthologous gene–SNP–trait trios, comparison of the distribution of colocalization posterior probability (PPH4) with human complex traits between trios that were significant versus not significant in cattle GTEx. P values were calculated using two-sided unpaired t tests. **g,** For the same 344 trios, comparison of the number of human complex traits showing evidence of colocalization between cattle GTEx–significant versus non-significant trios. P values were calculated using two-sided unpaired t tests. **h,** Example of a conserved whole-blood eQTL at *ZDHHC4* in cattle and humans that also influences human neutrophil count; human rs11559147 is homologous to the cattle ARS-UCD1.2 chr25_38409356 variant.

To further investigate whether cattle eQTL can provide evolutionary evidence for dissecting complex traits in humans, we employed S-LDSC to partition heritability of 133 human complex traits using orthologous variants of cattle eQTL^56^ (**Supplementary Table** 15). These orthologous variants were significantly enriched for heritability across all complex traits (**Supplementary Table** 16-17). Orthologous variants of tissue-specific cattle eQTL showed significantly higher enrichment than those of the remaining cattle eQTL, consistent with reports from human studies^57^ (**Fig.** 8e). For instance, compared to immune-specific eQTL detected in human GTEx, orthologous variants of cattle immune-specific eQTL displayed significantly higher enrichment in human immune-related traits (**Fig 8e; Extended Data Fig.** 10), supporting the idea that immune-related regulatory programs showed substantial conservation between cattle and humans^5,6^. Furthermore, we performed colocalization between human GTEx eQTL and 133 human GWAS traits, yielding 508,191 SNP–gene–tissue trios (PP.H4 > 0.8). Among them, 344 SNP–gene–tissue trios were tested in the CattleGTEx (**Extended Data Fig.** 10c**; Supplementary Table** 18), and 158 of them were also eQTL in cattle (FDR < 0.05), which were significantly more than that of MAF-matched random variants (*P*-value = 0.0021) (**Extended Data Fig.** 10d). Moreover, trios that were significant in the CattleGTEx exhibited stronger colocalization (higher PP.H4 values) with human complex traits than non-significant trios (*P*-value = 4.36×10^-13^; **Fig. 8f**) and were associated with more complex traits (**Fig. 8g**, *P*-value = 2.22×10^-2^). For instance, a variant (rs11559147 in humans and rs210096698 in cattle) showed conserved regulatory effects on *ZDHHC4* expression in whole blood in both humans and cattle, and it was significantly colocalized with a GWAS locus of neutrophil count in humans (PP.H4 = 0.99; **Fig. 8h**). The *ZDHHC* family encodes palmitoyltransferases that catalyze reversible protein S-palmitoylation^58^, thereby modulating membrane localization and signalling complex assembly of immune regulators and fine-tuning key immune signalling cascades^59,60^. Altogether, these results suggest the presence of evolutionary constraint regulatory variants between humans and cattle, which can facilitate the prioritization of causal variants and genes associated to complex traits and diseases in humans.

## Discussions

The CattleGTEx Phase1, built on 12,422 RNA-seq samples of 43 tissues from 8,956 animals worldwide, provided a comprehensive multi-tissue catalogue of regulatory effects of 4,317,531 genomic variants in cattle. Compared with the pilot phase, we extended the number of targeted tissues from 23 to 43, revealing more tissue-specific regulatory effects. In addition, reaching an average increase of 378 samples per tissue allow us to investigate non-primary variants. We also systematically explored pleiotropic regulatory effects of genomic variants on seven molecular phenotypes, revealing the distinct genetic architecture underlying molecular phenotypes, particularly between abundance and structure-based phenotypes. By examining selection signatures of *B. taurus* vs. *B. indicus* and dairy vs. beef, we demonstrated that both long-term adaptive evolution and short-term intensive artificial breeding were more likely to act on genomic variants with tissue-specific regulatory effects, which is also agreed with findings in sheep^15^. During divergence between *B. taurus* and *B. indicus*, selection signals were most pronounced in immune tissues, adipose, and skin, with enriched pathways related to immune responses such as bacterial invasion, consistent with adaptation of *B. indicus* to heat, humidity, and parasite exposure in tropical environments^2^. In contrast, dairy-beef divergence is not driven by environmental pressures but by strong artificial selection for production traits, including milk and meat yield^3^. Accordingly, selection signals in this comparison were concentrated in nervous and digestive tissues, with enriched pathways related to growth, metabolism, and production. This finding agreed with previous studies, highlighting the importance of the nervous system in responses to artificial selection for dairy cattle and associated milk production traits^18,61^.

The CattleGTEx Phase1 provided novel insights into the regulatory mechanisms underlying 44 complex traits in cattle. This enabled us to link 1,545 (75%) of GWAS loci to molecular phenotypes, genomic variants, target genes, and relevant tissues, representing a substantial improvement over the 260 (13%) colocalized GWAS loci reported in the pilot CattleGTEx study for the same complex traits. The different categories of molQTL exhibited distinct contributions to complex traits, with non-primary effects, pleiotropic eQTL, and tissue-specific molQTL showing higher enrichment for GWAS loci compared to the rest of regulatory variants. For the strong enrichment of non-primary and tissue-specific molQTL with GWAS loci, one possible explanation is that variants with large regulatory effects or effects shared across many tissues are more likely to perturb multiple biological processes and therefore incur stronger fitness costs. In contrast, regulatory variants that act in more restricted contexts, such as secondary regulatory layers or specific tissues, may avoid these strong selection pressures, allowing them to persist in populations and contribute to phenotypic variation detected by GWAS^62^. In addition, the enrichment between vertical pleiotropic eQTL and GWAS loci may reflect buffering mechanisms that operate across diverse biological processes or layers along the central dogma to main the stability of the whole system. Many genetic perturbations at a single molecular layer might be compensated through regulatory feedback, network redundancy, or post-transcriptional control. However, genetic variants that simultaneously affect multiple layers of molecular regulation may be more likely to bypass such buffering mechanisms. Their effects can therefore propagate more effectively through molecular networks and ultimately manifest at the organismal level, increasing the probability that they contribute to complex trait variation detected by GWAS. These findings also underscore the importance of using larger sample sizes to detect more variants with small, additive effects, profiling more molecular phenotypes, and considering additional tissue/cell types to better understand the genetic and molecular architecture underlying complex traits^30–33^. By aligning fine-mapped cattle eQTL to the human genome, we revealed evolutionary constraints on regulatory effects, facilitating the prioritization of causal variants and genes associated with complex traits and diseases in humans. For instance, a variant exhibited conserved regulatory effects on ZDHHC4 expression in blood in both humans and cattle and was significantly colocalized with neutrophil count in humans. Previous studies have reported that ZDHHC4 enhances the efficacy of immunotherapy, a role that has been experimentally validated^63^.

Despite these advances, we noted several limitations that will be addressed through future development of the CattleGTEx Project: **1**) only common SNPs were considered in this phase. More RNA-seq samples with paired WGS, improved reference genomes (i.e., telomere-to-telomere genomes^64^ and pangenomes^65^), and enhanced functional annotations (e.g., by integrating CAGE data to refine transcription start site resolution and mitigate tissue-specific annotation artifacts^66^) will enable us to explore regulatory effects of rare variants, transposable elements, short tandem repeats and complex structural variants. **2**) The current dataset remains biased toward commercial breeds, limiting the representation of global genetic diversity in cattle. Future efforts should prioritize the inclusion of local and indigenous breeds worldwide, which are shaped by distinct ecological and environmental conditions, to better capture regulatory mechanisms underlying local environmental adaptation and conservation-relevant traits^67^. **3**) This study focused on molecular phenotypes that were derived from short-read RNA-seq. Long-read RNA-seq data will be essential for more accurate quantification of molecular phenotypes, particularly isoform expression and alternative splicing^68^. Moreover, integrating additional molecular layers, such as profiles from epigenomics, proteomics, metabolomics, and gene regulatory networks^69–72^, will be essential to capture downstream and complex regulatory mechanisms that cannot be resolved at the transcriptomic level alone. **4**) We only conducted molQTL mapping at bulk-tissue resolution and therefore did not fully capture regulatory heterogeneity across cell types and states^73–75^. Although we explored cell-type-dependent regulation using computational deconvolution approaches in a companion study^21^, population-scale single-cell molQTL mapping^73,76^, together with spatially resolved transcriptomic data^77,78^, will be required to dissect regulatory mechanisms operating at finer cellular and spatiotemporal scales. **5**) This study considered a limited set of biological contexts, primarily focusing on different tissue types and breeds. Future development will incorporate additional contexts, including sex, developmental stages, and diverse environmental conditions^79–81^, which will require the establishment of a FarmGTEx tissue biobank with standardized sampling protocols, controlled environmental conditions, and comprehensive metadata collection, to characterize context-dependent regulatory mechanisms in cattle and further advance our understanding of the genetic architecture of complex traits. **6**) The regulatory variants prioritized through computational analyses in this study remain largely inferential. Experimental validation, including *in vitro* assays and *in vivo* functional studies^82,83^, will be crucial to confirm causal regulatory effects and to translate these findings into mechanistic understanding of complex traits.

## Author contribution statement

L.Fang conceived and designed the project. H.Li., H.Zhang and Z.Wei performed bioinformatic analyses of RNA-Seq data analysis. P.Zhao conducted whole-genome sequence data analysis. H.Zhang performed multi-omics analysis. H.Li and H.Zhang conducted molQTL mapping. H.Li performed GWAS integrative analysis. H.Li and P.Zhao performed selection region analysis. D.Zhu performed comparative analysis between human and cattle. L.Fang., H.Li., S.Peter., Y.Zhou., X.Wang., X.Lu., Q.Zhang., J.Li., Y.Wu., W.Li., J.Cole., Z.Cai., G.Sahana., D.Liu., Y.Li., D.Sun., Y.Yu., Y.Zhang., B.Han., H.Sun., M.Jia., S.Zhu., X.Liu., P.Su., L.Jiang., Y.Liu., W.Lyu., E.Clark., M. Littlejohn., R.Xiang., Y.Chen., Z.Pan., Y.Hou., M.Li., C.Xu., Q.Liu., Q.Yang., Z.Yang., H.Jiang., X.Lu., M. Wijst., E. Mármol-Sánchez., H.Zhou., L.Guan., X.Zhao., I. Ionita-Laza., J.Grady., D.MacHugn., L.Frantz., A.Leonard., H.Pausch., O.Madsen., M.Gòdia., M.Derks., R.Crooijmans., E.Ibeagha-Awemu., M.Wang., Y.Wang., W.Liu., S.Chen., J.Tian., M.Tian., H.Zhang., B.Lin., B.An., C.Fang., J.Du., Z.Zhang., S.Zhao., Y.Gao., Y.Tang., Q.Lin., J.Teng., D.Guan., X.Liu., W.Zheng., Q.Zhang and M.Gong contributed to the critical interpretation of analytical results before and during manuscript preparation. Y.Wang., J.Sheng and M.Wang built the Cattle-GTEx Phase1 web portal. L.Fang., M.Lund., B.Li., Y.Wang and J.Jiang. contributed to the data and computational resources. H.Li, H.Zhang, D.Zhu., and L.Fang drafted the manuscript. All authors read, edited, and approved the final manuscript.

## Online Methods

### Ethics and sample collection

The vast majority of RNA-seq samples and all WGS are publicly available. No Ethics of animal sampling are not applicable. For the newly generated data, all animal procedures and experimental protocols were approved by the Institutional Animal Care and Use Committee of Northwest A&F University following the recommendations of the Regulations for the Administration of Affairs Concerning Experimental Animals of China.

### Bulk RNA-seq data processing

We downloaded 24,975 RNA-seq datasets from the Sequence Read Archive (SRA, https://www.ncbi.nlm.nih.gov/sra) and 1,385 datasets from the Genome Sequence Archive (GSA, https://ngdc.cncb.ac.cn/gsa) using the Aspera command-line tool (**Supplementary Table** 1-2). In addition, we newly generated 1,268 RNA-seq datasets from 3 tissues of 533 Holstein cattle, including 514 samples in lung, 459 sample in rumen and 295 samples in lymph node (See **Supplementary Notes**). After raw RNA sequencing data were obtained, the adapters were removed along with low quality reads using Fastp (v0.23.4) with parameters: --trim_front 3, --trim_tail 3, --length_required 36, --cut_right, --cut_window_size 4 and --cut_mean_quality 15^84^. Clean reads were then aligned to the B. taurus reference genome (ARS-UCD1.2, Ensembl v108^85^) using STAR (v 2.7.10b) with parameters: --chimSegmentMin 10, --twopassMode Basic, --outFilterMismatchNmax 10, -- chimOutJunctionFormat 1, --outFilterType BySJout, --alignSJoverhangMin 8 and -- alignSJDBover-hangMin 1^86^. In total, 19,790 samples with >10M mapped reads, >40% uniquely mapped reads, and >40% expressed genes were retained for downstream analyses. The detailed mapping quality of all the RNA-seq samples is summarized in **Supplementary Table** 4.

### Genotype calling and imputation of RNA-seq samples

PCR duplicates were removed in bam files produced by STAR aligner previously using the MarkDuplicates module of Genome Analysis Toolkit (GATK, v4.3.0.0)^87^, and then SplitNCigarReads module was engaged to split reads that contained Ns in their cigar string. Next, GATK BaseRecalibrator module and ApplyBQSR module were applied to recalibrate base quality score based on the Ensembl dbSNP database (v108). The recalibrated BAM files were used for direct genotype imputation with GLIMPSE2 (v2.0.0)^88^, based on a multi-breed genotype reference panel^31^. Variants with minor allele frequency (MAF) < 0.05 or imputation quality score (INFO) < 0.75 were subsequently filtered out using BCFtools (v1.17)^89^, resulting in a total of 4,317,531 SNPs. After obtaining the genotype files, WASP (v0.3.4) was used to remove mapping bias of RNA-seq data^90^. SNPs from each chromosome were extracted separately using BCFtools (v1.17)^89^, and the snp2h5 script provided in the WASP package was used to convert the resulting VCF files into HDF5 format^90^. These HDF5 files were subsequently used to correct mapping bias for each sample. The find_intersecting_snps.py script was applied to extract reads overlapping with SNPs, and STAR (v2.7.10b) aligner was then employed to remap these reads using the same parameters as in the initial mapping^86^. Reads for which one or more allelic versions failed to remap to the same genomic location were filtered out using filter_remapped_reads.py. Finally, duplicate reads were removed using rmdup.py or rmdup_pe.py for single- end and paired-end RNA-seq data, respectively.

### Quantification of molecular phenotypes

After mapping bias removal, the corrected bam file was used for molecular phenotype definition and quantification. Normalized expression (TPM) of 23,103 protein-coding and lncRNA genes (Bos_taurus. ARS-UCD1.2, Ensembl v108) were generated using Stringtie (v.2.2.1)^91^. Raw read counts for all 20,875 genes and 102,033 exons were obtained with featureCounts (v2.0.3)^92^. Transcribed enhancer regions were annotated, and a custom SAF file was prepared after excluding those overlapping with gene bodies^93^, then the raw counts were also quantified using featureCounts (v.2.0.3)^92^. For transcript-level expression, Salmon (v1.10.1)^94^ was used in mapping-based mode with clean reads as input to quantify transcript expression, generating both TPM and raw count outputs. The index for Salmon was built with the decoy sequences generated using SalmonTools^94^. For alternative splicing, Leafcutter (v.0.2.9)^95^ was used to quantify the splicing events as following: 1) mapping bias–corrected BAM files from the previous step were converted into junction files using the bam2junc.sh script provided by Leafcutter; (2) intron clusters were identified with leafcutter_cluster.py and annotated to genes using map_clusters_to_genes.R; (3) introns with zero read counts in more than 50% of samples or with fewer than max(10 or 0.1n) unique values in each tissue (where n is the sample size) were removed; (4) low-complexity introns were filtered out based on the z score of cluster read fractions across individuals; (5) intron excision ratios were calculated, and the filtered counts were converted to BED format using prepare_phenotype_table.py. Finally, excision levels of introns were normalized as percent spliced-in (PSI) values. For the quantification of 3’UTR alternative polyadenylation (3’UTR APA), Dapars (v2.1) was used with the following steps^96^: 1) distal polyadenylation sites from the Ensembl annotation (v108) were extracted using the script DaPars_Extract_Anno.py provided by Dapars (v2.1) package, 2) genomecov function in BEDTools (v2.31.0) was utilized to obtain wiggle alignment files^97^, 3) all the wiggle files were utilized to quantify APA usage within each tissue^98,99^. The output represents the percentage of distal polyA site usage index (PDUI) for each gene in each sample. Loci with more than 50% missing values were excluded, and the remaining entries were imputed using the K nearest neighbor (KNN) method^22,100^. For RNA stability estimation, genomic coordinates of exon and intron were extracted, and HTSeq was used to quantify raw exon and intron read counts^101,102^. The constitutive exon-to-intron ratio was then calculated for each gene. Genes with more than 50% missing values were excluded, and the remaining missing entries were imputed using the KNN approach^22^. Allele specific expression analysis was conducted using phASER (v1.1.1)^103^. Imputed genotype VCF file together with bam file from WASP software were input to phASER tool for phasing variants with options of “--mapq 2 --baseq 10”^103^. Then, gene-level haplotype counts were obtained using the script phaser_gene_ae.py. Then all the individual haplotypic counts file were aggregated by tissue using the script phaser_expr_matrix.py.

### molQTL mapping

Within each tissue, duplicated samples (IBS > 0.9) were removed. The molQTL mapping was performed using a linear mixed model implemented in OmiGA (v1.0.3)^37^. SNPs with MAF < 0.05 or minor allele count (MAC) < 6 were excluded with “--mac-threshold 6 --maf-threshold 0.05” in each of tissue. We defined the *cis*-window of genes as ±1 Mb of TSS and obtained the nominal *P* values of *cis*-molQTL with the option of “--mode cis”, while selecting the suitable phenotypic PCs as co-variables using the optional of “--dprop-pc-covar 0.001”^14,37^. Covariables with high co-linearity were removed with the option of “--rm-collinear-covar 0.95”. We then employed two layers of multiple testing corrections using OmiGA^37^. In the first layer, we applied an adaptive permutation approach to calculate the empirical *P* values of variants within each gene and obtained the permutation *P* value of the lead variant for each gene. In the second layer, we conducted the clipper correction for the nominal *P* values and computed the FDR of lead variants across all tested genes using the option of “--multiple-testing clipper”^104^. A gene was considered as an molGene if at least one SNP had a corrected *P* < 0.05 and a *P*-value below the gene-level permutation-derived threshold calculated by OmiGA (v1.0.3)^37^; and molQTL were defined as SNPs with nominal P values below the gene-level significance threshold.

### Conditional and fine-mapping analysis of molQTL

To detect multiple independent *cis*-molQTL signals of each molGene, we employed two complementary approaches. First, conditionally independent molQTL were identified using a stepwise regression procedure implemented in OmiGA (v1.0.1.3) with “--mode cis_independent”, which iteratively identifies the most significant SNPs while conditioning on previously selected variants^37^. We further applied “Sum of Single Effects” (SuSiE) model (v.1.0)^105^ to fine-map causal QTL and define credible sets of variants, which uses a Bayesian multiple regression framework to estimate the posterior inclusion probability (PIP) for each variant. Variants with PIP ≥ 0.8 were considered putative causal variants, and 95% credible sets were constructed to capture the most likely causal variants within each signal.

### Colocalization analysis between molQTL

To investigate whether molecular phenotypes share genetic regulatory mechanisms with eQTL, we conducted colocalization analysis between each molecular phenotype with eQTL. Colocalization was performed using the *coloc.abf* function in the *coloc* (v5.2.3) R package^106^, which implements an approximate Bayes factor framework to identify shared causal variants. For each pair of molecular phenotypes, the method computes posterior probabilities for five hypotheses: (H0) no association with either trait; (H1) association only with the molecular phenotype; (H2) association only with the QTL; (H3) association with both traits but with independent causal variants; and (H4) association with both traits and a shared causal variant. As the results, we defined the colocalized signals between two molQTL by PP.H4 > 0.8, whereas the distinct signals by PP.H3 < 0.25^21^. In addition, LD between the lead SNPs of colocalized signals was calculated using PLINK (v1.9) to further characterize shared regulatory loci^107^.

### Tissue-sharing/specificity analysis of molQTL

We used multivariate adaptive shrinkage (MashR, v0.2.57)^108^ to characterize cross-tissue sharing patterns of molQTL. For the overall estimation, we extracted the z scores (slope / slope_se) of the top molQTL for each molGene across all tissues in which it was detected and used these top-effect molQTL–gene pairs as input to MashR. In addition, one million molQTL-gene pairs were randomly sampled from all nominal associations tested by OmiGA across tissues^37^, and their corresponding z-scores were obtained. Pairs with missing values in more than five tissues were removed to avoid inflating tissue-specific signals driven by sparse observations rather than true biological effects; for the remaining pairs, missing z scores were assigned a standard error of 1×10⁶. MashR was then applied to estimate the local false sign rate (LFSR) for each tissue, and molQTL with LFSR < 0.05 were considered active in that tissue. Tissue similarity was assessed by computing pairwise Spearman correlations of effect sizes among active molQTL.

### Functional enrichment of molQTL

To assess whether molQTL were enriched in functional regions, we obtained functional annotations for all tested SNPs using SnpEff (v5.2)^109^ and, separately, chromatin-state annotations from the Cattle FAANG project^55^. Enrichment analyses were performed independently for each annotation type. For each tissue and each annotation type, a 2×2 contingency table was constructed comparing (i) SNPs identified as molQTL and (ii) SNPs carrying the given annotation. Odds ratios (ORs), 95% confidence intervals, and *P*-values were computed for each QTL annotation pair. ORs and confidence intervals were derived from the 2×2 tables, with continuity adjustments applied when necessary. *P*-values were obtained using a chi square test of independence. In each tissue, all SNPs tested by OmiGA were used as the background for enrichment calculations.

### GWAS summary statistics of complex traits

To investigate the regulatory mechanisms underlying complex traits in cattle, we systematically integrated identified eQTL with GWAS summary statistics for 44 complex traits in Holstein cattle^110^, including 22 body conformation, 6 milk production, 8 reproduction, and 8 health traits. Independent SNPs were identified for each trait using the *GCTA-cojo* function in GCTA (v1.94.1)^111^, based on the Holstein population in the reference genotype imputation panel^31^. For colocalization analysis, independent SNPs with *p*-values less than 1 × 10⁻⁵ were then selected as suggestive significant, and corresponding QTL regions were defined by extracting all SNPs located within ±1 Mb of each lead SNP^21^.

### Enrichment analysis between *cis*-molQTL and GWAS loci

QTLenrich (v2.0), the software assesses enrichment of common variant associations for a given complex trait among molQTL, was used for enrichment analysis between our seven molQTL and GWAS loci from 44 complex traits^54^. Three input files were prepared, 1). Top pairs were generated with lead SNPs of molGenes in each tissue; 2). Significant pairs were generated with SNPs in which the nominal *P*-value larger than the permutation *P*-value in each gene; 3). All pairs were generated with all SNPs used in molQTL mapping. All parameters were set as default. We next performed multiple testing by using p.adjust function in R (v4.0.2) with the parameter *Method = “BH”* across P-values from all 1,540 tissue-trait pairs and calculated FDR values^112^. Significant enrichment results (FDR < 0.05) were used for statistical plot.

### *cis*-molQTL-GWAS colocalization

To identify shared genetic variants between GWAS loci and molQTL, we conducted a colocalization analysis between each molGene and overlapped GWAS loci using the *coloc.abf* function in the *coloc* package (v5.2.3)^106^, which is an approximate Bayes factor colocalization analysis. The package computed posterior probabilities for (1) no association with either GWAS loci and molQTL (*H*0); (2) association only with GWAS loci (*H*1); (3) association only with molQTL (*H*2); (4) association with both GWAS loci and molQTL but two independent signals (*H*3); and (5) association with both molecular phenotype and shared signals (*H*4). The colocalized signals between GWAS loci and molQTL were defined by PP.H4 > 0.8^21,22^. Moreover, we calculated the LD of two lead SNPs for a pair of GWAS variants and molQTL using PLINK (v.1.9)^107^.

### Function enrichment analysis of gene lists

Gene Ontology (GO) and Kyoto Encyclopedia of Genes and Genomes (KEGG) analyses were performed using the clusterProfiler (v.4.0)^113^. The GO terms and KEGG pathways of selected genes were enriched in the org.Bt.eg.db and KEGG-bta databases, using the erichGO and enrichKEGG functions, respectively, with a threshold parameter of “pvalueCutoff = 0.05”. We considered a GO term or a KEGG pathway significant if it had FDR < 0.05.

### Population-stratified molQTL analysis

To investigate the difference in *cis*-eQTL between breeds, global ancestry of all samples within each tissue was estimated by ADMIXTURE (v1.3.0) with K=2^114^. To ensure clear genetic assignments of samples to four breed groups (i.e., *B. taurus, B. indicus,* dairy, and beef cattle), we matched the population labels with ancestral lineage composition, and considered the label correct if the lineage purity was greater than 80%. Subsequently, other samples with ancestral purity greater than 80% (even if the original label was not) were defined as belonging to this breed. We then conducted *cis*-eQTL mapping separately in *B. taurus*, *B. indicus*, dairy and beef cattle populations using the same procedures as described in **molQTL mapping** above. Replications of *cis*-eQTL between populations within a tissue and tissues within a population were quantified using Storey’s π_1_ statistic and Pearson’s correlation of normalized eQTL effect sizes and compared to replication between tissues within a population and populations within a tissue^22^. To further characterize sharing patterns across populations, we conducted a meta-analysis of *cis*-eQTL across tissues and population groups using the MashR package (v0.2.57)^108^, following the same pipeline used for **Tissue-sharing/specificity analysis of molQTL**.

### Selective sweep and molQTL enrichment analysis

To detect selection signatures between *B. taurus* and *B. indicus,* we selected 200 *B. taurus* (50 Holstein, 50 Jersey, 50 Simental and 50 Angus) and 142 *B. indicus* (20 Leiqiong, 20 Butana, 11 Horro, 11 Boran, 10 Arsi, 10 Goffa, 10 Mursi, 9 Barka, 9 Fogera, 9 Afar, 9 Sheko, 7 Baoule and 7 Nelore) with WGS data (>10x) from the genotype imputation panel^31^ (**Supplementary Table 3**) with 28,958,211 SNPs. To detect selection signatures between beef and dairy cattle, we selected 651 dairy (Holstein) and 552 beef (326 Simmental, 64 Angus, 60 Charolais, 58 Limousin and 44 Hereford) cattle from the genotype imputation panel^31^ (**Supplementary Table 4**) with 28,958,211 SNPs. We analyzed genetic differentiation (*F*_ST_) using the vcftools (v0.1.16) tool^115^, employing a sliding window of 30 Kb with a step size of 10 Kb. The windows were divided into deciles based on their *F*_ST_ values, forming ten groups (*F*_ST_1–*F*_ST_10) with increasing selection intensity. Enrichment analysis was then performed between each group and the molQTL from credible sets derived through fine mapping. Odds ratio (OR) was calculated to evaluate enrichment levels between selected regions and molQTL with the following formula^21^:

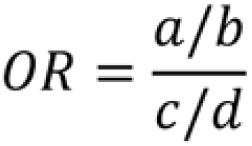

In which *a* represents the numbers of molQTL located in genomic regions of each *F*_ST_ groups, *b* represents the number of molQTL, *c* represents the number of SNPs located in genomic regions of each *F*_ST_ group, *d* represents the number of all SNPs. The frequency of alleles in each SNP was calculated using the option of “--freq” in Plink (v1.9)^107^.

### Cross-species conservation analysis between cattle and humans

To evaluate cross-species conservation between cattle and humans, we mapped fine-mapped cattle eQTL to the human reference genome (hg38) using the UCSC liftOver tool^116^. SNPs that were successfully converted were considered as orthologous (sequence-level conserved) variants between cattle and humans. To assess conservation at the functional level, we tested whether enrichment of eQTL within regulatory elements was preserved between species. Specifically, we performed: (i) enrichment analysis of cattle eQTL in cattle regulatory elements^55^; (ii) enrichment analysis of liftOvered cattle eQTL in human regulatory elements^117^; and (iii) enrichment analysis of human eQTL from GTEx v8 in human regulatory elements. We then quantified concordance in enrichment patterns across the three analyses using Pearson’s correlation to assess cross-species functional conservation.

In addition, we used the *AlphaGenome* model with *in silico* mutagenesis to predict the regulatory effects of orthologous variants of cattle eQTL in the human genome^53^. Only the tissue-level gene expression track in the *AlphaGenome* was considered in this study. Predicted effects were then compared across different eQTL groups and MAF-matched random SNPs.

We next collected GWAS summary statistics for 133 human complex traits (**Supplementary Table 7**) from the Pan-UK Biobank^118^ and applied LD score regression (*LDSC* v1.0.1)^56^ to estimate the proportion of SNP-heritability attributable to each predefined eQTL group. We conducted colocalization analysis between human GTEx v8 eQTL and GWAS of human complex traits using coloc v5.2.3^91^, and considered a posterior probability threshold of PP.H4 > 0.8 as high-confidence shared signals. To ensure biologically interpretable cross-species comparisons, we retained only tissue-gene-SNP trios that satisfied all of the following criteria: (i) genes have identifiable human– cattle orthologs; (ii) the tissue is matched between humans and cattle; and (iii) the orthologous locus is segregating in both the human GTEx and cattle GTEx datasets. As a control, we randomly selected an equal number of trios. We then tested whether SNPs from the retained trios were also associated with gene expression in CattleGTEx using a single-SNP linear mixed model, in which the genomic relationship matrix (GRM) was included as a random effect to account for population structure and relatedness; this analysis was performed using GCTA (v1.93.2)^111^. Fixed-effect covariates were kept identical to those used in the cattle eQTL mapping. SNPs with FDR < 0.05 were considered significant.

### Statistics and reproducibility

No statistical method was used to predetermine the sample size. The details of data exclusions for each specific analysis are available in the Methods section. For all the boxplots, the horizontal lines inside the boxes show the medians. Box bounds show the lower quartile (Q1, the 25th percentile) and the upper quartile (Q3, the 75th percentile). Whiskers are minima (Q1 − 1.5× IQR) and maxima (Q3 + 1.5× IQR), where IQR is the interquartile range (Q3–Q1). Outliers are shown in the boxplots unless otherwise stated. The experiments were not randomized, as all the datasets are publicly available from observational studies. The investigators were not blinded to allocation during experiments and outcome assessment, as the data were not from controlled randomized studies.

## Data availability

All raw data analyzed in this study are publicly available for download without restrictions from SRA (https://www.ncbi.nlm.nih.gov/sra/) and GSA (https://ngdc.cncb.ac.cn/gsa). The ARS-UCD1.2 (v.110) cattle reference genome is available at Ensembl (https://www.ensembl.org). Details of RNA-Seq samples, WGS samples, molQTL summary results, selection analysis results, GWAS information of human disseases, colocalization, TWAS and SMR results can be found in Supplementary Tables, respectively. All processed data and the full summary statistics of molQTL mapping and genotype imputation reference panel are available at https://cattlegtex.kiz.ac.cn/.

## Code availability

All the computational scripts and codes for RNA-Seq and WGS analyses, as well as the ene_expression_analyses, molQTL mapping, fine mapping, conditional analysis, tissue sharing analysis, functional enrichment, colocalization, TWAS, SMR, selection signal analysis and LDSC are available at the FarmGTEx GitHub website (https://github.com/FarmGTEx/CattleGTEx_Phase1_Pipeline_v0).

## Acknowledgments

1. L. F. was supported by Agriculture and Food Research Initiative Competitive grants nos. 2022-67015-36215 from the USDA National Institute of Food and Agriculture, seed-funding from CellFood Hub (AUFF), Danmarks Frie Forskningsfond nos. 5254-00069B. B. L. acknowledges funding from the UK Biotechnology and Biological Sciences Research Council (BBSRC) with grant BB/X009505/1.

---

**Extended Data Fig. 1.**
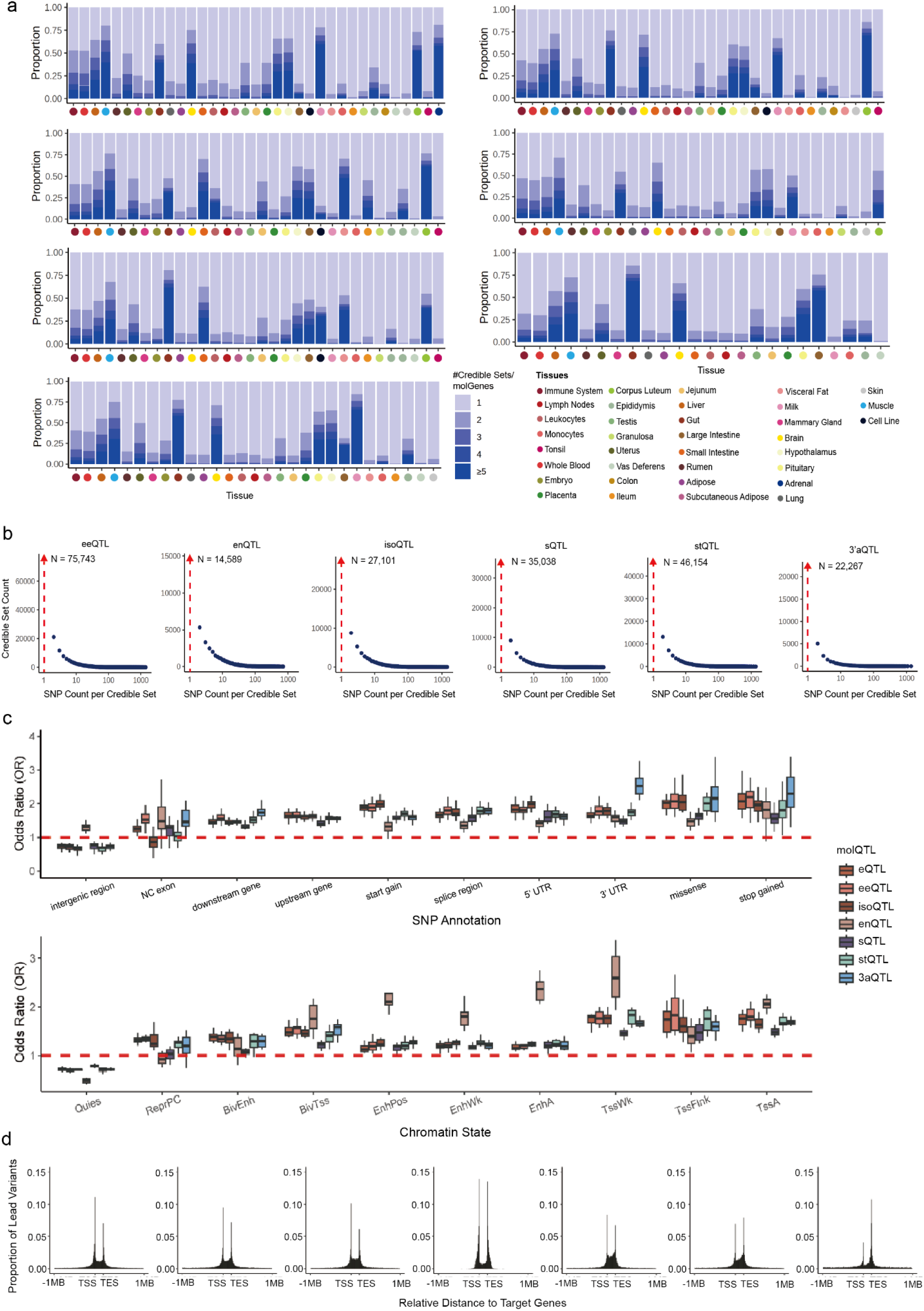
**a,** The distribution of the number of fine-mapped *cis*-molQTL per gene across seven molQTL is shown as blue stacked bars across 35 tissues. Tissues are ordered by decreasing sample size. **b,** The size distribution of fine-mapped credible sets for all identified *cis*-molQTL across 35 tissues. The red arrow indicates count of credible sets containing only one single candidate variant. **c,** Functional enrichment of lead SNPs in seven molQTL with regulatory elements (top: SNP classification, bottom: chromatin states). **d,** Distribution of the distance to TSS regions across lead SNPs in seven molQTL.

**Extended Data Fig. 2.**
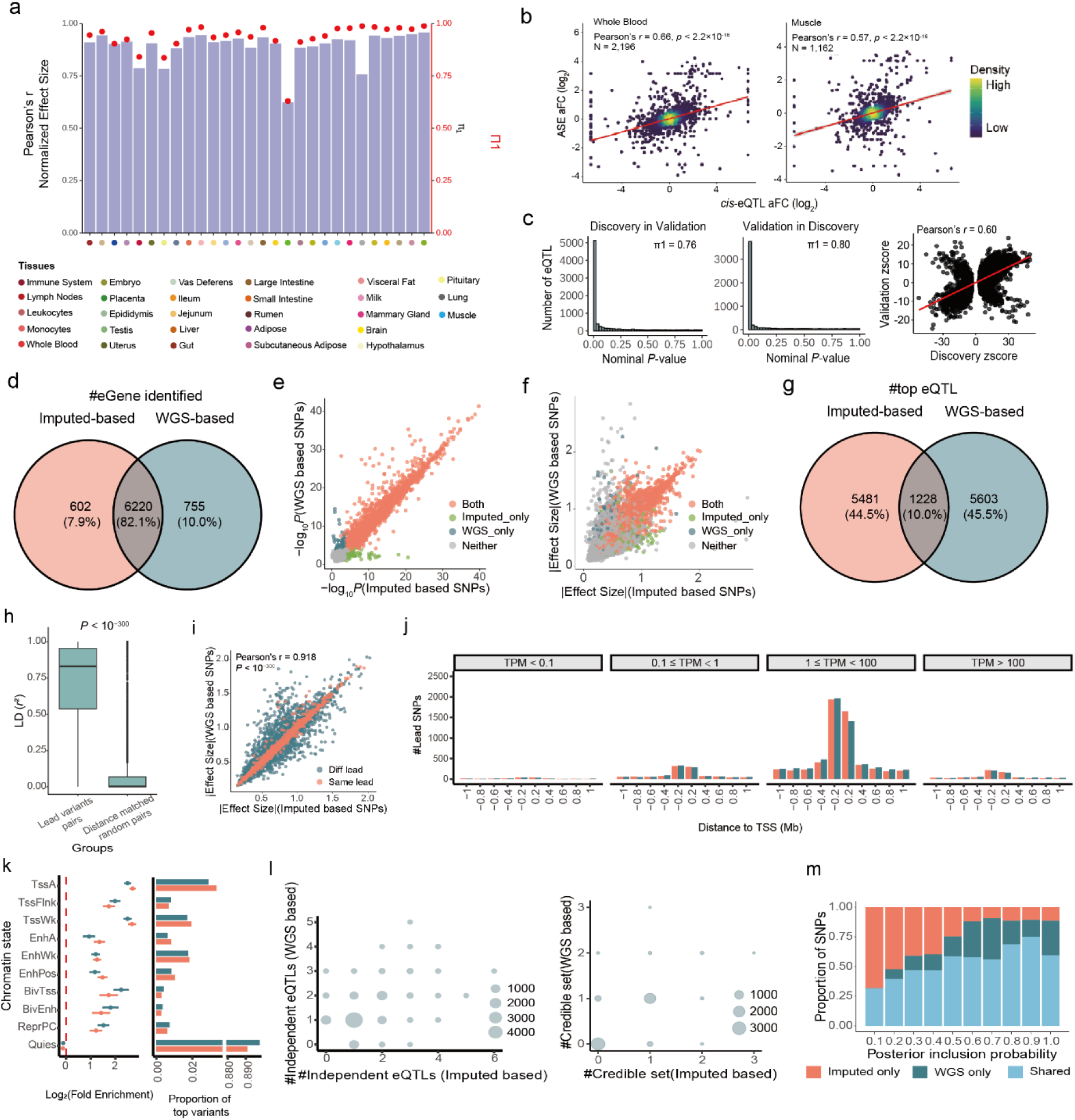
**a,** Internal validation of *cis*-eQTL. We selected 28 tissues with sample sizes > 100 and randomly split samples into equally sized discovery and validation cohorts. eQTL mapping was performed independently in each cohort. Bars show Pearson’s correlation of normalized effect sizes between the two groups, and red dots represent Storey’s π_1_ statistics, indicating the replication rate of *cis*-eQTL. **b,** Spearman’s correlation between *cis*-eQTL effect sizes (allelic fold change, log₂ scale) and corresponding effect sizes derived from allele-specific expression (ASE) analysis. **c,** External validation of *cis*-eQTL using an independent Holstein population (*n* = 99). Storey’s π_1_ statistics and Spearman’s correlation were used to assess replication between external population and CattleG-TEx Phase1 population in blood. **d,** Robustness of our approach by comparing molQTL mapping using RNA-seq imputed versus whole-genome sequencing (WGS)-called genotypes in an independent blood-tissue cohort (n = 99). Venne plot represents the overlapping of eGenes detected using imputed versus WGS–called genotypes. **e,** Comparison of significance (-log10 P) across different gene categories. Each point represents a gene. Genes are categorized as Both (n = 5,798): eGene detected by both WGS–called and imputed genotypes; WGS_based_only (n = 552): eGene detected only by WGS–called genotypes; Imputed_based _only (n = 1,792): eGene detected only by imputed genotypes; Neither (n = 7,935): Non-eGene in either dataset. **f,** Comparison of effect sizes (beta) of top variants across different gene categories. Each point represents a gene. **g,** Overlap of top variants using imputed versus WGS–called genotypes. **h,** Linkage disequilibrium (LD, r2) among lead variants of the same eGenes (n = 5,142) identified using imputed- and WGS-called. The “Distance-matched random set” contains SNP pairs with similar genomic distances as the lead variant pairs. The boxplot shows the median as the central line, the 25th–75th percentiles as the box limits, and whiskers extending 1.5× the interquartile range. Statistical significance was determined using a two-sided Student’s t-test. **i,** Correlation of effect sizes for lead variants of eGenes identified using imputed- and WGS-based genotypes. Same lead indicates that the shared eGenes have the same lead variant, while Diff lead indicates that the shared eGenes have distinct lead variants. P-values were calculated using the asymptotic t approximation. **j,** Number of top variants detected using imputed and WGS-called genotypes, plotted by distance to the TSS and stratified by median gene expression across samples. **k,** Functional enrichment (log2 Fold change, mean ± s.d., left) and proportion (right) of eQTL detected using WGS-called versus imputed genotypes across chromatin states. **l,** The left panel represents numbers of independent eQTL per gene detected using WGS-called and imputed genotypes. Point size reflects the number of eGenes with the corresponding number of independent eQTL. The right panel represents numbers of credible sets per gene detected in fine-mapping using WGS-called and imputed genotypes. Point size reflects the number of eGenes with the corresponding number of credible sets. **m,** Percentage of fine-mapped variants detected using WGS-called and imputed genotypes, shown across different posterior inclusion probability (PIP) thresholds

**Extended Data Fig. 3.**
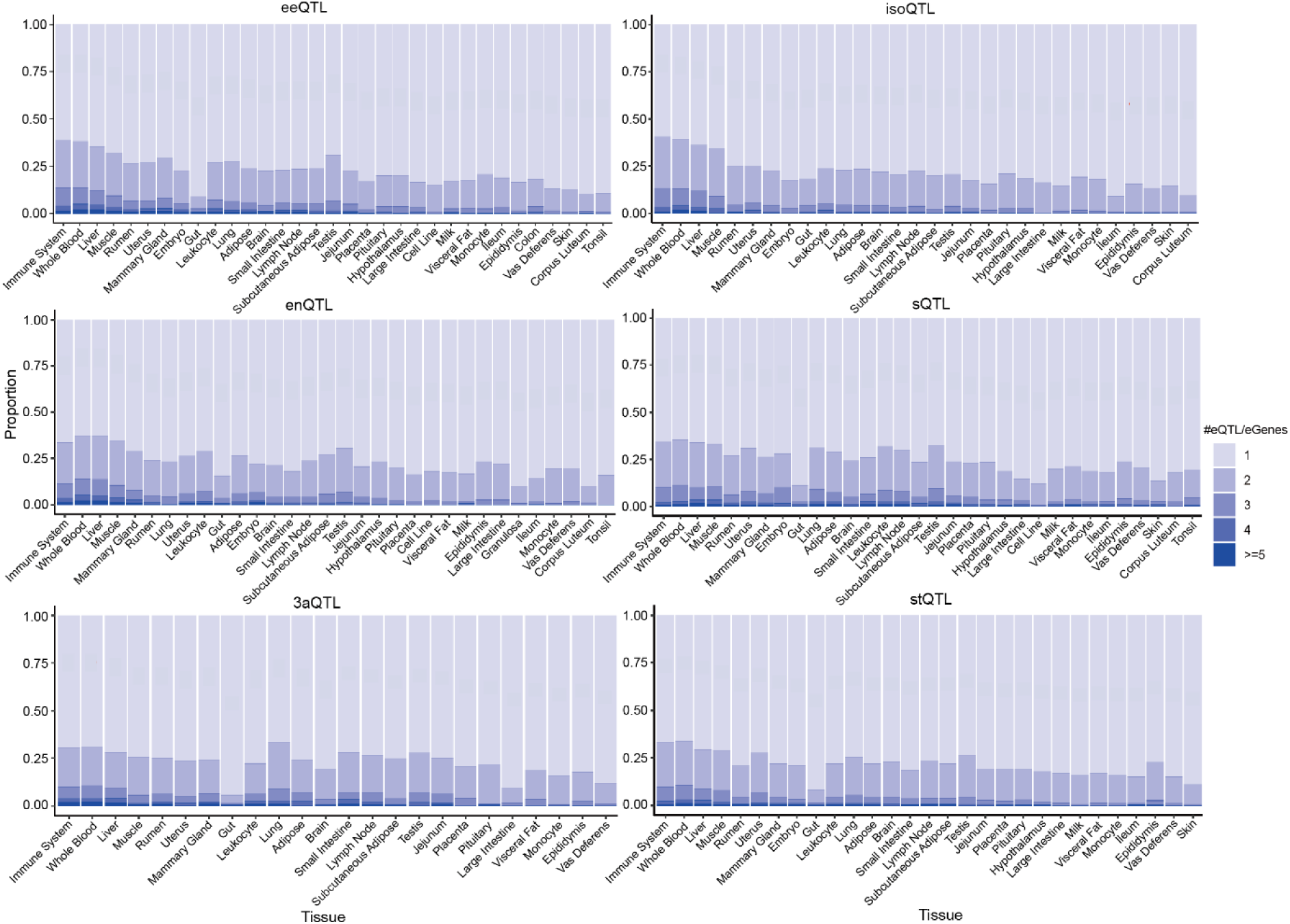
The distribution of the number of conditionally independent *cis*-molQTL per gene across six molQTL is shown as blue stacked bars across 35 tissues. Tissues are ordered by decreasing sample size.

**Extended Data Fig. 4.**
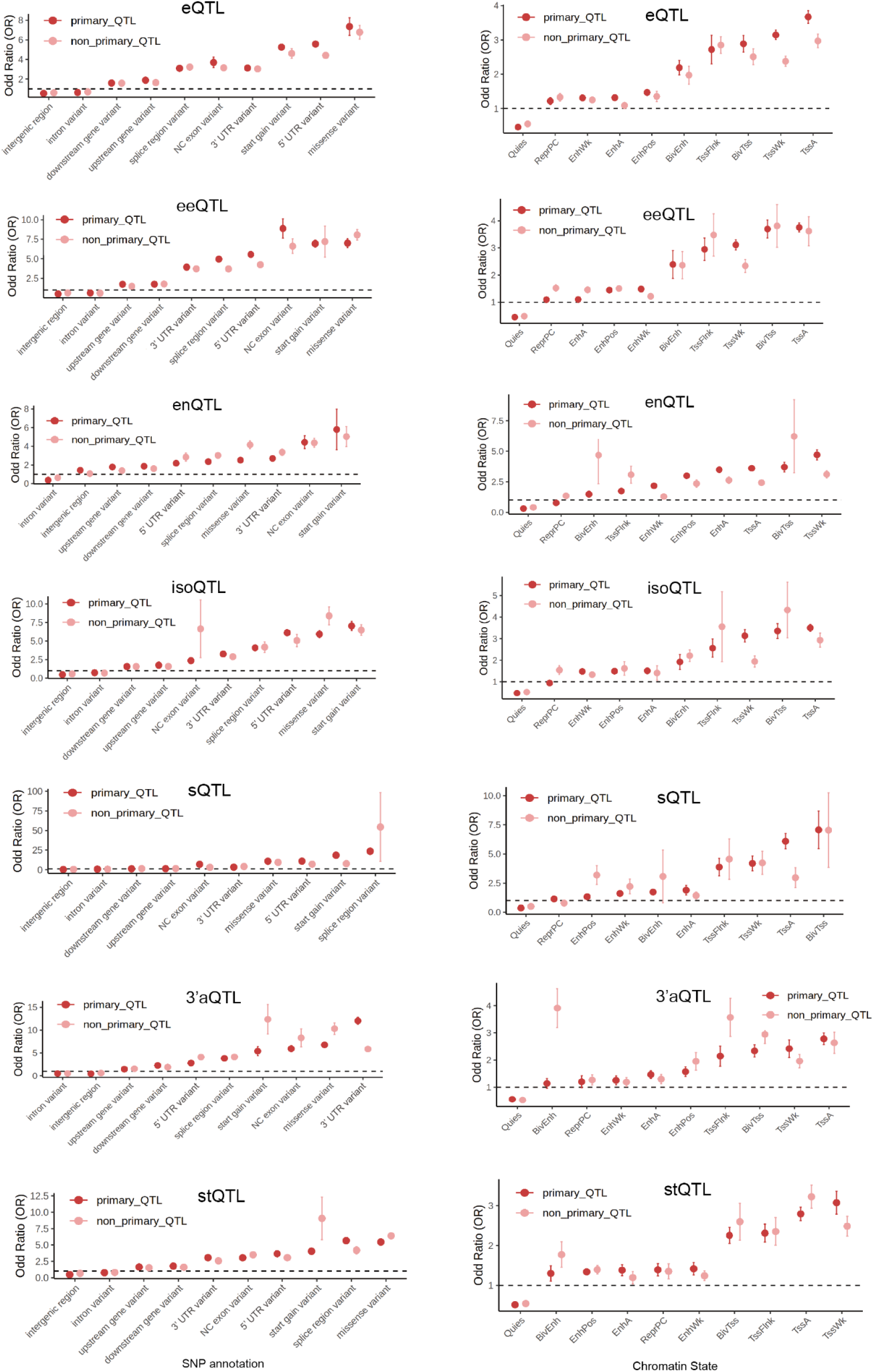
Functional enrichment of primary and non-primary effects in six molQTL with regulatory elements (left: SNP classification, right: chromatin states).

**Extended Data Fig. 5.**
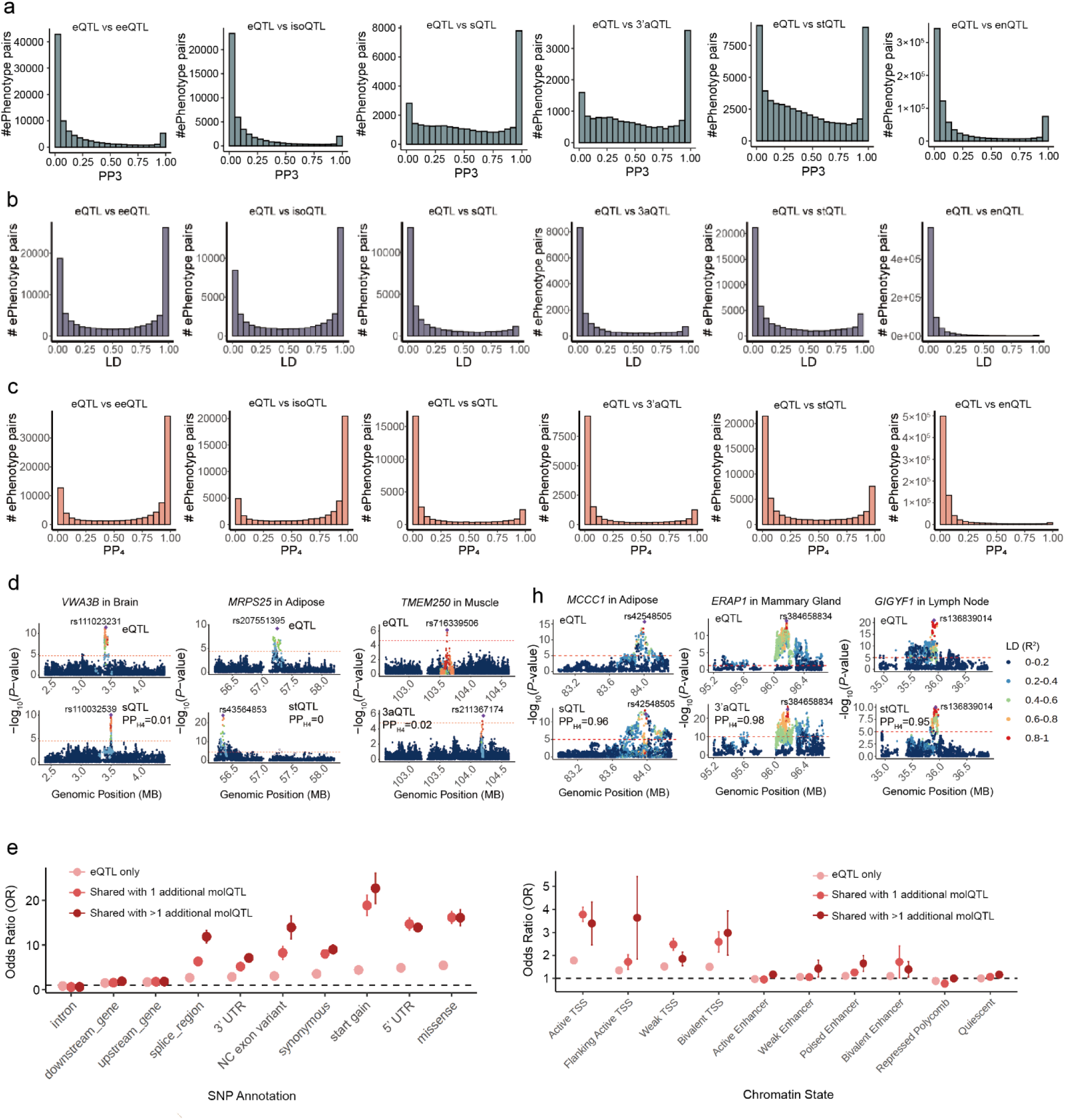
**a-c,** Colocalization analyses eQTL across other six molecular phenotypes of the same molGene. PP_3_: the posterior probability that the two molQTL are driven by distinct causal variants (**a**). PP_4_: the posterior probability that the two molQTL share a single causal variant (**c**). LD (r^2^): the linkage disequilibrium of lead SNPs of two molecular phenotypes derived from the molGene (**b**). **d,** Examples of distinct signals between eQTL and other six molQTL within the same molGene. The two-sided P values of lead SNPs were calculated by the linear mixed regression model. **e,** Functional enrichment of pleiotropic eQTL with regulatory elements (left: SNP classification, right: chromatin states).

**Extended Data Fig. 6.**
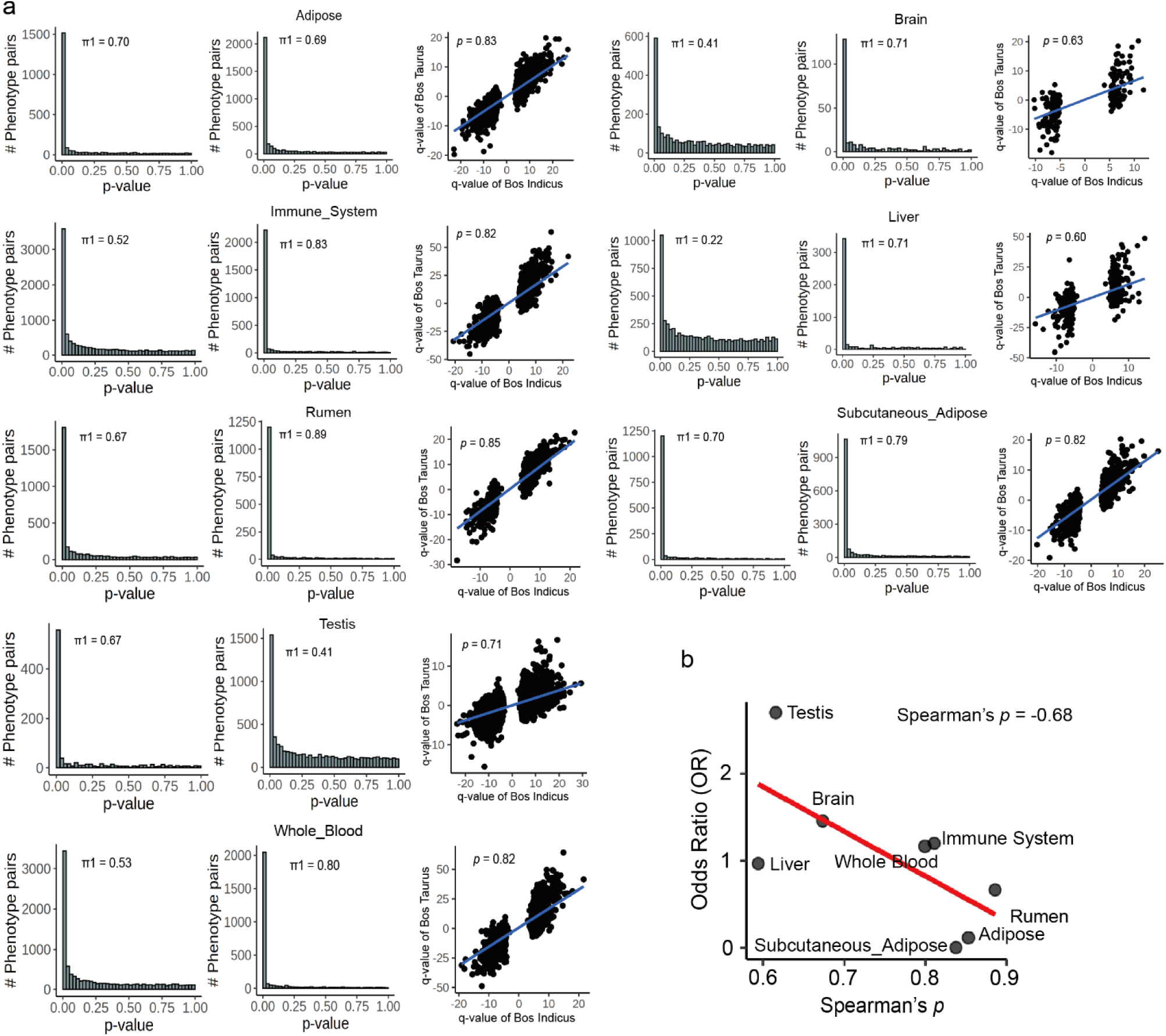
**a,** Storey’s π_1_ statistics implemented in the qvalue package (v2.32.0)^119^ and Spearman’s correlation were used to assess replication of effect sizes between *B. taurus* and *B. indicus* across eight tissues. In each tissue, the p-value of the eQTL of *B. taurus* was used as observation (**left**) and validation data (**mid**), respectively. **b,** Spearman’s correlation between odds ratio (OR) of enrichment between tissue-specific eQTL and strongly selected regions and spearman’s correlation of effect sizes between *B. taurus* and *B. indicus*.

**Extended Data Fig. 7.**
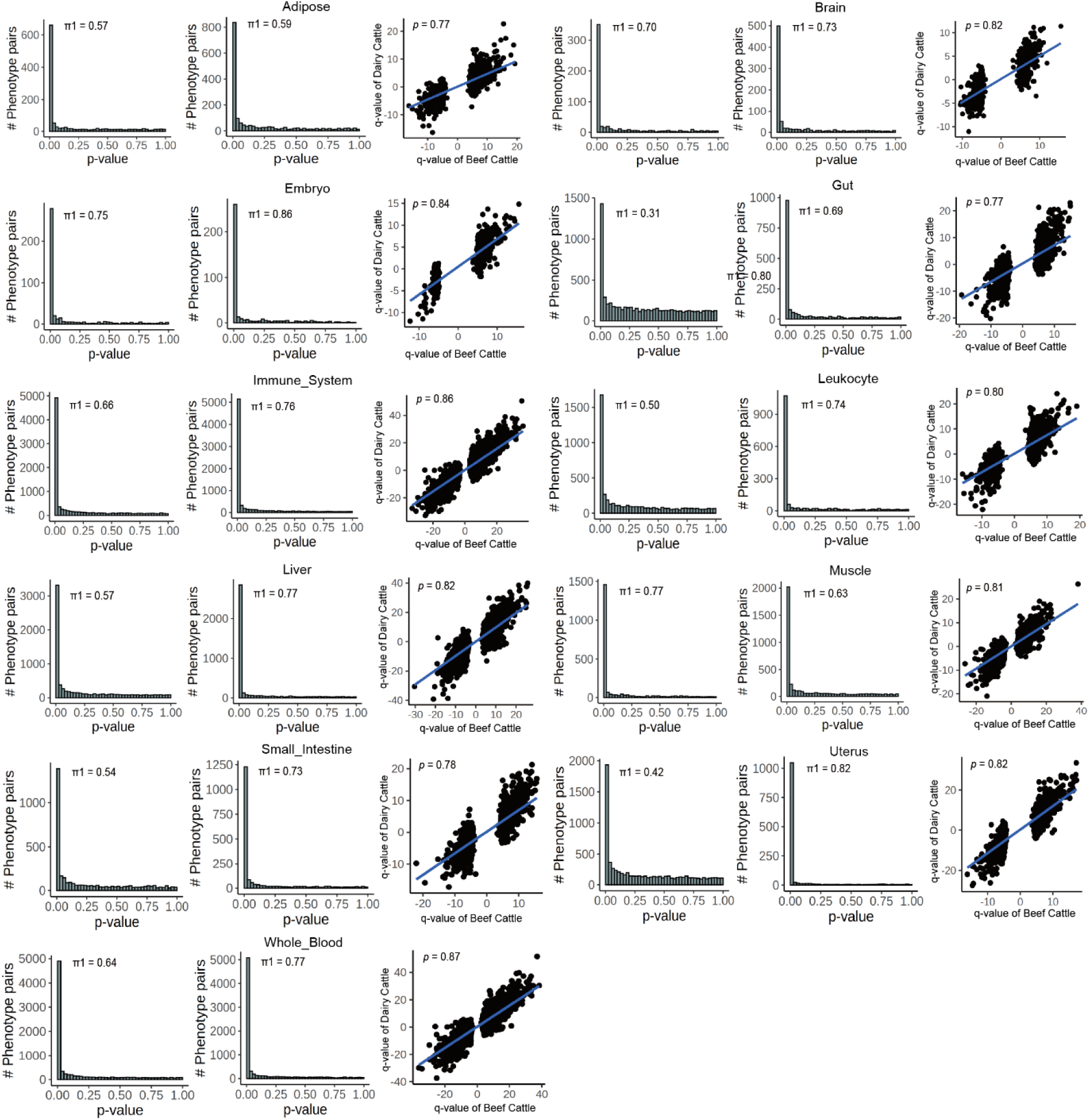
Storey’s π_1_ statistics implemented in the qvalue package (v2.32.0)^119^ and Spearman’s correlation were used to assess replication of effect sizes between dairy cattle and beef cattle across 11 tissues. In each tissue, the p-value of the eQTL of dairy cattle was used as observation (**left**) and validation data (**mid**), respectively.

**Extended Data Fig. 8.**
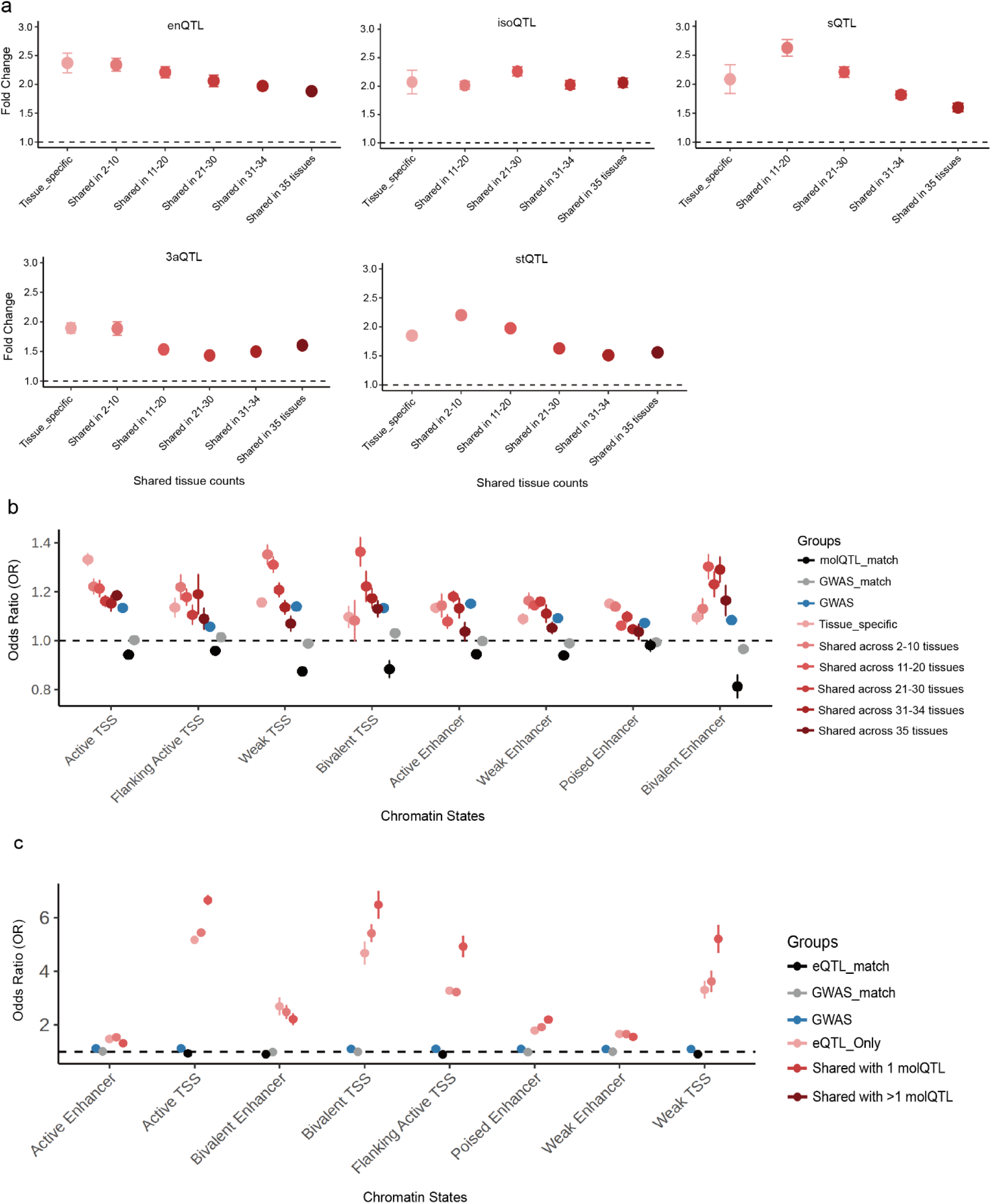
**a,** Enrichment (odds ratio) of tissue specific/shared molQTL with genomewide associations (GWAS) of 44 cattle traits using QTLenrich (v.2.0)^54^. **b,** Functional enrichment between tissue specific/shared molQTL and regulatory elements. **c,** Functional enrichment between pleiotropic eQTL and regulatory elements.

**Extended Data Fig. 9.**
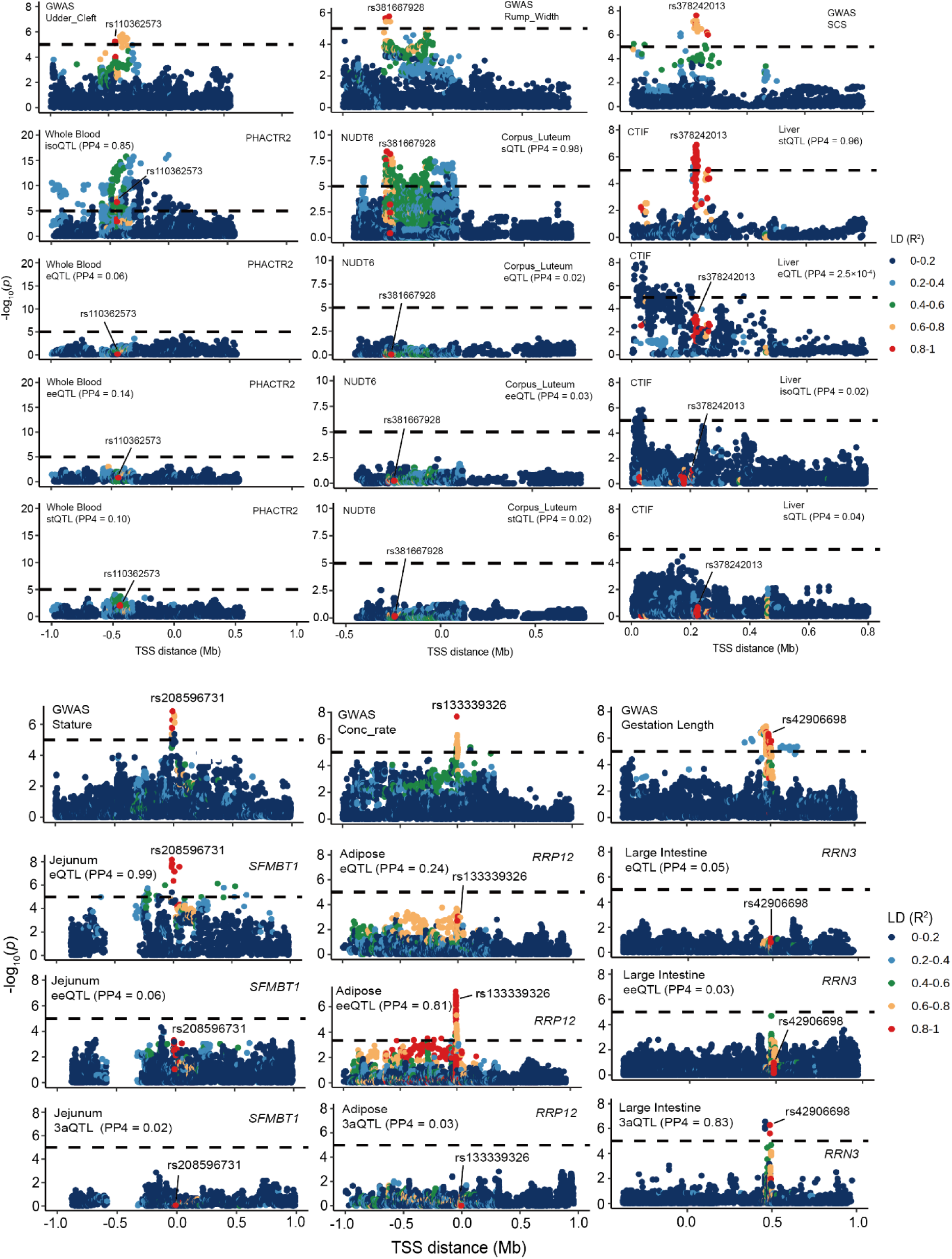
Manhattan plots showing the -log₁₀(p) distribution of distinct colocalization between GWAS loci and each of seven molQTL, including udder cleft (GWAS) and isoQTL in whole blood across genetic variants within the locus of *PHACTR2*, rump width (GWAS) and sQTL in corpus leteum across genetic variants within the locus of *NUDT6*, SCS (GWAS) and stQTL in liver across genetic variants within the locus of *CTIF,* stature (GWAS) and eQTL in jejunum across genetic variants within the locus of *SFMBT1,* conception rates (GWAS) and eeQTL in adipose across genetic variants within the locus of *RRP12* and gestation length (GWAS) and 3’aQTL in large intestine across genetic variants within the locus of *RRN3*. Variants are colored according to their linkage disequilibrium (LD, r²) with the lead SNP. The two-sided P values of SNPs were calculated by the linear mixed regression model.

**Extended Data Fig. 10.**
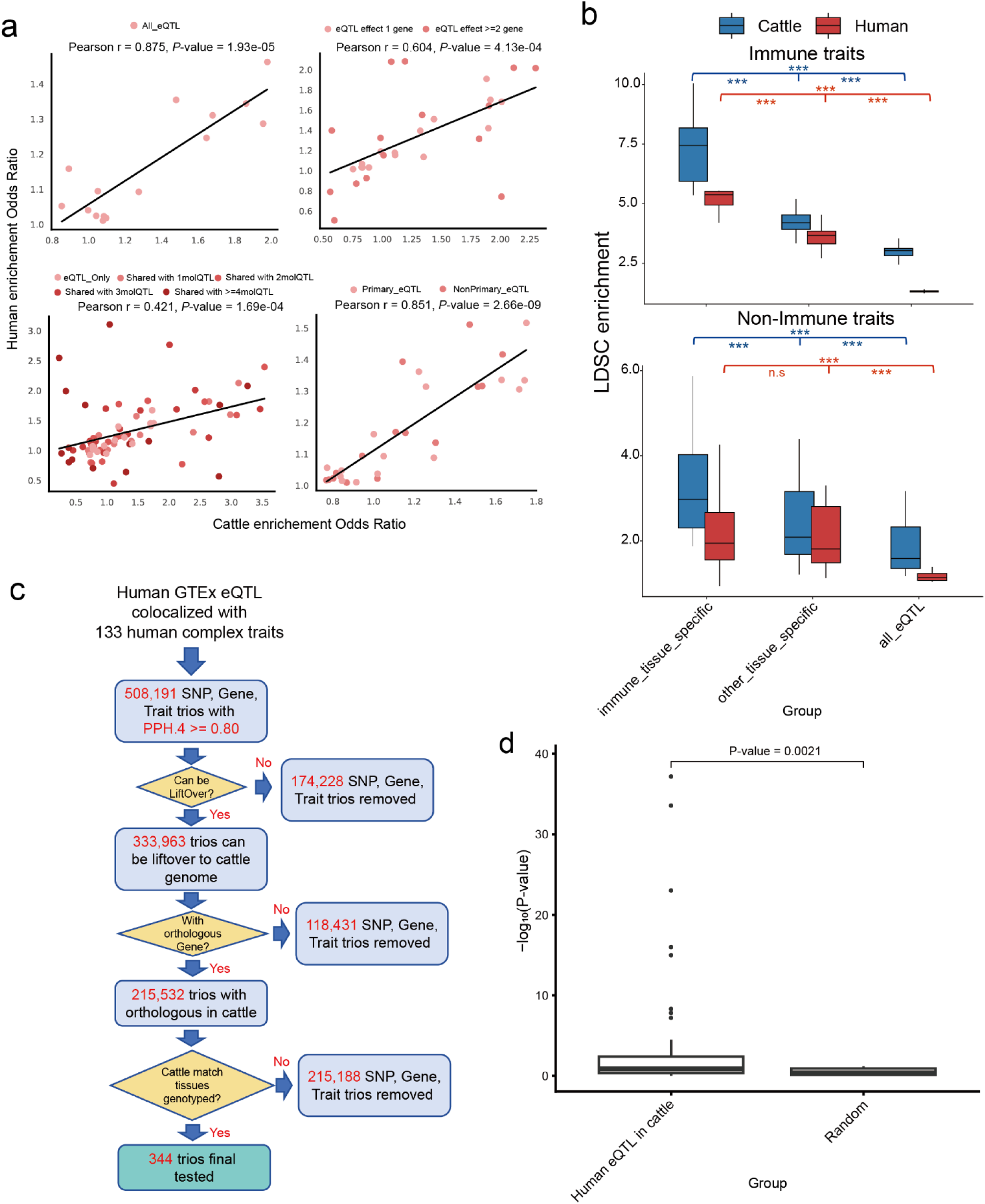
**a,** Correlation between enrichment odds ratios (ORs) for different cattle eQTL groups in cattle FAANG annotations and the corresponding enrichment ORs after liftOver to the human genome in ENCODE annotations. **b,** Comparison of stratified LD score regression (S-LDSC) heritability enrichment of tissue-specific eQTL annotations from cattle and humans across immune versus non-immune traits. P values were calculated using two-sided unpaired t tests; P < 0.05 (), P < 0.01 (), and P < 0.001 (). **c,** Schematic of filtering steps and counts used to derive cattle orthologous, segregating trios from 508,191 gene–variant–trait trios obtained by colocalization between human GTEx eQTL and 133 human complex traits. Filters were applied sequentially: (1) the human variant could be lifted over from the human genome to the cattle genome; (2) the corresponding eGene had an ortholog in cattle; (3) the corresponding tissue was available in the cattle dataset; and (4) the lifted-over cattle locus harbored a segregating SNP. This yielded 344 retained trios. **d,** Comparison of association significance in cattle for the 344 retained trios tested using a single-variant linear mixed model versus a random set of matched orthologous loci. P values were calculated using two-sided unpaired t tests.

